# How short decoding times, stimulus dimensionality and spontaneous activity constrain the shape of tuning curves: A speed-accuracy trade-off

**DOI:** 10.1101/2022.09.09.505677

**Authors:** Movitz Lenninger, Mikael Skoglund, Pawel Herman, Arvind Kumar

## Abstract

According to the efficient coding hypothesis, sensory neurons are adapted to provide maximal information about the environment given some biophysical constraints. Early sensory neurons modulate their average firing rates in response to some features of the external stimulus, creating tuned responses. In early visual areas, these modulations (or tunings) are predominantly single-peaked. However, periodic tuning, as exhibited by grid cells, has been linked to a significant increase in decoding performance. Does this imply that the tuning curves in early visual areas are sub-optimal? We argue that the time scale at which neurons encode information is imperative to understanding the relative advantages of single-peaked and periodic tuning curves. Because, if decoding ability scales differently with time for the different shapes of tuning curves, the time scale at which the neurons operate becomes critical. Here, we show that the possibility of catastrophic (large) errors due to overlapping neural responses for distinct stimulus conditions creates a trade-off between decoding time and decoding ability. Unfortunately, standard theoretical measures such as Fisher information do not capture these errors. We investigate how (very) short decoding times and stimulus dimensionality affect the optimal shape of tuning curves for stimuli with finite domains. In particular, we focus on the spatial periods of the tuning curves (or the number of “peaks”) for a class of circular tuning curves. We show a general trend for minimal decoding time, i.e., the shortest decoding time required to produce a statistically reliable signal, to increase with increasing Fisher information implying a trade-off between accuracy and speed. This trade-off is reinforced whenever the stimulus dimensionality is high or there is ongoing activity. Thus, given constraints on processing speed, we present normative arguments for the existence of single-peaked, rather than a periodic, tuning organization observed in early visual areas.

## Introduction

One of the fundamental problems in systems neuroscience is understanding how sensory information can be represented in the spiking activity of an ensemble of neurons. The problem is exacerbated by the fact that individual neurons are highly noisy and variable in their responses, even to identical stimuli (Arieli et al., 1996). A common feature of early sensory representation is that the neocortical neurons in primary sensory areas change their average responses only to a small range of features of the sensory stimulus. For instance, some neurons in the primary visual cortex respond to moving bars oriented at specific angles (Hubel and Wiesel, 1962). This observation has led to the notion of *tuning curves*. Together, a collection of tuning curves provides a possible basis for a neural code.

A considerable emphasis has been put on understanding how the structure of noise and correlations affect stimulus representation given a set of tuning curves (Shamir and Sompolinsky, 2004; Averbeck and Lee, 2006; Franke et al., 2016; Zylberberg et al., 2016; Moreno-Bote et al., 2014; Kohn et al., 2016). More recently, the issue of local and catastrophic errors, which dates back to the work of Shannon (Shannon, 1949), has been raised in the context of neuroscience (Xie, 2002; Sreenivasan and Fiete, 2011). Intuitively, local errors are small estimation errors that depend on the trial-by-trial variability of the neural responses and the local shapes of the tuning curves surrounding the true stimulus condition (Fig. 1a, see *s*_1_). On the other hand, catastrophic errors are very large estimation errors that depend on the trial-by-trial variability and the global shape of the tuning curves (Fig. 1a, see *s*_2_). While a significant effort has been put into studying how stimulus tuning and different noise structures affect local errors, less is known about the interactions with catastrophic errors. For example, *Fisher information* is a common measure of the accuracy of a neural code by providing a measure of the sensitivity to local stimulus changes in the output responses of a given population (Brunel and Nadal, 1998; Abbott and Dayan, 1999; Guigon, 2003; Moreno-Bote et al., 2014; Benichoux et al., 2017). Intuitively, if Fisher information is high, small changes in the stimulus conditions produce large changes in the population’s spike count statistics. The use of Fisher information is justified by the Cramér-Rao bound, by which it can be related to the minimal attainable mean squared error (MSE) for any unbiased estimator. However, because Fisher information can only capture local errors, the true MSE of a system might be considerably larger in the presence of catastrophic errors (Xie, 2002; Kostal et al., 2015), especially if the available decoding time is short (Bethge et al., 2002; Finkelstein et al., 2018).

**Figure 1:**
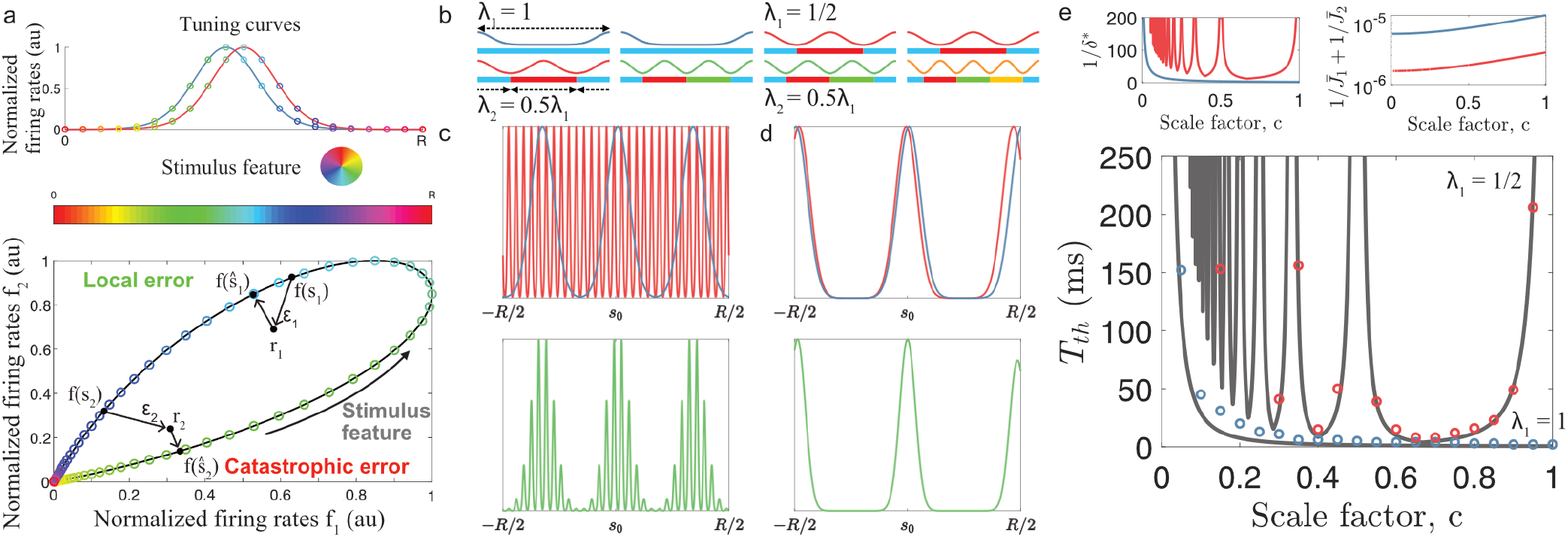
Minimal decoding times in two module populations. a) (top) A two-neuron system encoding a single variable using tuning curves. (bottom) The tuning curves create a one-dimensional activity trajectory embedded in a two-dimensional neural activity space (black trajectory). Decoding the two stimulus conditions, *s*_1_ and *s*_2_, illustrates the two types of estimation errors that can occur due to trial-by-trial variability, local 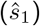 and catastrophic 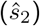. b) Illustration of the individual likelihood functions formed by two-module systems. Below the likelihood functions, the stimulus interval has been partitioned and color-coded according to the spatial period of the tuning curves. Note that a single module cannot differentiate stimulus conditions across these partitions, i.e., the stimulus is only unambiguous within each color-coded spatial period. c) Using a second module with a spatial period much smaller than the first module can introduce stimulus ambiguity both within the correct mode of the first module’s likelihood and across the first module’s modes. In this illustration, the individual likelihoods have been perturbed just enough around the correct stimulus *s*_0_ such that the joint likelihood function is ambiguous. Note that the perturbations can be very small, especially if *c* ≫ 1. d) Same as in c) but for spatial periods that are similar but not identical. In this case, catastrophic estimation errors only occur due to modes of the joint likelihood function far from the true stimulus condition becoming larger than for the correct mode. e) The dependence of the scale factor c on the minimal decoding time. The blue and red circles indicate the simulated minimal decoding times for populations with λ_1_ = 1 and λ_1_ = 1/2, respectively. The gray lines indicate the estimation of the minimal decoding times according to Eq. 8, with *p_error_* = 10^−3^. The insets show 1) the predicted value of 1/*δ*\*, and 2) the inverse of the Fisher information. Note that the color codes correspond to the color code of the circles in the main figure. See Table S1 in Methods for list of parameter values.

A curious observation is that the tuning curves of early visual areas predominately use single firing fields, whereas grid cells in the entorhinal cortex are known for their periodically distributed firing fields (Hafting et al., 2005). That is, in the early visual areas tuning curves are mostly single-peaked while grid cells have numerous, periodically distributed peaks. The multiple firing locations of grid cells increase the precision of the neural code compared to single-peaked tuning curves (Sreenivasan and Fiete, 2011; Mathis et al., 2012; Wei et al., 2015). This raises the question of why periodic firing fields are not a prominent organization of early visual processing, too?

The theoretical arguments in favor of periodic tuning curves have mostly focused on local errors under the assumption that catastrophic errors are negligible (e.g., Sreenivasan and Fiete (2011)). However, given the response variability, it takes a finite amount of time to accumulate a sufficient number of spikes to decode the stimulus. Given that fast processing speed is a common feature of visual processing (Thorpe et al., 1996; Fabre-Thorpe et al., 2001; Rolls and Tovee, 1994; Resulaj et al., 2018), it is crucial that each neural population in the processing chain can quickly produce a reliable stimulus-evoked signal. Therefore, the time required to produce signals without catastrophic errors will likely put fundamental constraints on any neural code, especially in early visual areas.

Here, we contrast Fisher information with the minimal decoding time required to remove catastrophic errors (i.e., before Fisher information becomes a valid descriptor of the MSE). We base the results on the maximum likelihood estimator for uniformly distributed stimuli (i.e., the maximum a posteriori estimator) using populations of tuning curves with an inhomogeneous number of firing fields (i.e., the number of “peaks”). We show that minimal required decoding time tends to increase with increasing Fisher information. This suggests a trade-off between the accuracy of a neural signal and the speed by which it can be reliably produced. Furthermore, we show that the difference in minimal decoding time grows with the number of jointly encoded stimulus features and in the presence of spontaneous activity. Thus, single-peaked tuning curves require shorter decoding times and are more robust to spontaneous activity than periodic tuning curves. Finally, we exemplify the issue of large estimation errors and periodic tuning in simple spiking neural networks tracking either a step-like stimulus change or a continuously time-varying stimulus.

## Results

### Shapes of tuning curves, Fisher information and catastrophic errors

To enable a comparison between single-peaked and periodic (multi-peaked) tuning curves, we consider circular tuning curves responding to a D-dimensional stimulus **s** ∈ [0, *R*)^*D*^ according to

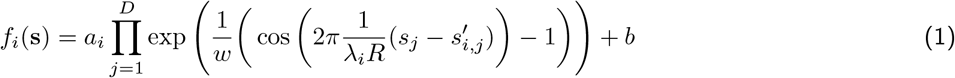

where *a_i_* is the peak amplitude of the stimulus-related tuning curve *i, w* is a width scaling parameter, λ*_i_* defines the spatial period (relative to the stimulus range, *R*) of neuron *i*, *s_i,j_* determines the location of the firing field(s) in the *j*:th dimension, *D* is the number of stimulus dimensions, and *b* determines the amount of spontaneous activity. The parameters are kept fixed for each neuron, thus ignoring any effect of learning or plasticity. In simulations, the stimulus domain was set to [0,1)^*D*^ for simplicity, although this choice does not qualitatively affect the results.

Throughout this paper, we assume that the stimulus is uniformly distributed and that its dimensions are independent of each other. This can be seen as a worst-case scenario as it maximizes the entropy of the stimulus. In a single trial, we assume that the number of emitted spikes for each neuron is conditionally independent and follows a Poisson distribution given some stimulus-dependent rate *f_i_*(*s*). Thus, the probability of observing a particular activity pattern given the stimulus-dependent rates is

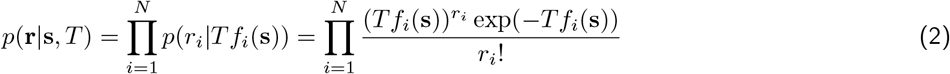

where **s** denotes the stimulus vector, *N* the number of neurons in the population, and **r** a vector of spike counts (of length *N*).

Given a model of neural responses, e.g., Eq. 2, the Cramér-Rao bound provides a lower bound on the accuracy by which the population can communicate a signal. The Cramér-Rao bound states that a lower limit of the MSE for any unbiased estimator is given by the inverse of Fisher information (Lehmann and Casella, 1998). Thus, increasing Fisher information reduces the lower bound on the minimal attainable MSE. For this reason, Fisher information has been a popular measure for studying population coding. For sufficiently large populations, using the population and spike count models in Eq. 1 and Eq. 2, Fisher information is given by (see Sreenivasan and Fiete (2011) or Methods for details)

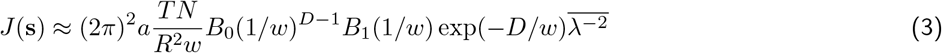

where 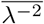 denotes the average of the squared inverse of the (relative) spatial periods across the population, and *B_α_*(·) denotes the modified Bessel function of the first kind. Eq. 3 (and similar expressions) suggests that populations consisting of periodic tuning curves, for which 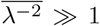, are superior at communicating a stimulus signal than a population using tuning curves with only single firing fields, where 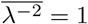. However, (inverse) Fisher information only predicts the amount of local errors for an efficient estimator. Hence, the presence of catastrophic errors (Fig. 1a) can be identified by large deviations from the predicted MSE for an asymptotically efficient estimator. Therefore, we defined minimal decoding time as the shortest time required to reach the Cramér-Rao bound.

### Periodic tuning curves and stimulus ambiguity

To understand why the minimal decoding time can differ with different spatial periods, consider first the problem of stimulus ambiguity that can arise with periodic tuning curves. If all tuning curves in the population share the same relative spatial period, λ, then the stimulus-evoked responses can only provide unambiguous information about the stimulus in the range [0, λ*R*). Beyond this range, each stimulus condition has a statistically identical response distribution for a stimulus condition λ*R* away. Thus, single-peaked tuning curves (λ = 1) provide unambiguous information about the stimulus (Fig. 1b). Periodic tuning curves (λ < 1), on the other hand, require the use of tuning curves with two or more distinct spatial periods to resolve the stimulus ambiguity (Fiete et al., 2008; Mathis et al., 2012; Wei et al., 2015). In the following, we assume the tuning curves are organized into discrete modules, where all tuning curves within a module share a spatial period (Fig. 1b) mimicking the organization of grid cells (Stensola et al., 2012). For convenience, assume that λ_1_ > λ_2_ > … > λ_*L*_ where *L* is the number of modules. Thus, the first module provides the most coarse-grained resolution of the stimulus interval, and each successive module provides an increasingly fine-grained resolution. It has been suggested that a geometric progression of spatial periods, such that λ_*i*_ = *c*λ_*i*–1_ for some spatial factor 0 < *c* ≤ 1, may be optimal for maximizing the resolution of the stimulus while reducing the required number of neurons (Mathis et al., 2012; Wei et al., 2015). However, trial-by-trial variability can still cause stimulus ambiguity and catastrophic errors - at least for short decoding times, as we show later, even when using multiple modules with different spatial periods.

### Minimal decoding times in two module populations

How does the choice of spatial periods impact the minimal decoding time? To get some intuition, we first consider the case of using only two different spatial scales. From the perspective of a probabilistic decoder (Seung and Sompolinsky, 1993; Deneve et al., 1999; Ma et al., 2006), assuming that the stimulus is uniformly distributed, the maximum likelihood (ML) estimator is Bayesian optimal (and asymptotically efficient). The maximum likelihood estimator aims at finding the stimulus condition which is the most likely cause of the observed activity, **r**, or

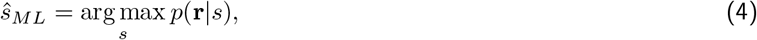

where *p*(**r**|*s*) is called the likelihood function. The likelihood function equals the probability of observing the observed neural activity, **r**, assuming that the stimulus condition was *s*. In the case of independent Poisson spike counts (or at least independence across modules), each module contributes to the joint likelihood function *p*(**r**|*s*) with individual likelihood functions, *Q*_1_ and *Q*_2_ (Wei et al., 2015). Thus, the joint likelihood function can be seen as the product of the two individual likelihood functions, where each likelihood is λ*_i_R*-periodic

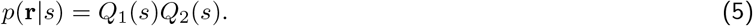

In this sense, each module provides its own ML-estimate of the stimulus, 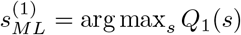 and 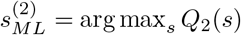, but because of the periodicity of the tuning curves there can be multiple modes for each of the likelihoods (e.g., Fig 1c and d, top panels). For the largest mode of the joint likelihood function to be also centered close to the true stimulus condition, the distance between 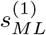 and 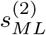 must be smaller than between any other pair of modes of *Q*_1_ and *Q*_2_. We assume that the stimulus estimate from each module is efficient, that is, the difference *δ* in the modes closest to the true stimulus *s*_0_ is a normally distributed random variable

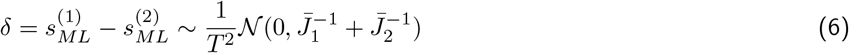

where 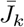 refers to the time-normalized Fisher information of module *k*. Thus, as the decoding time *T* increases, the variance in the distances between the “true” modes of *Q*_1_ and *Q*_2_ decreases. Hence, it is necessary for the decoding time *T* to be large enough such that it is rare for *δ* not to be the smallest distance between any two modes of *Q*_1_(*s*) and *Q*_2_(*s*).

To limit the probability of the decoder experiencing catastrophic errors to some small error probability *p_error_*, we impose that

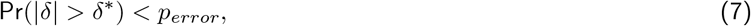

where *δ\** is the largest allowed distance of *δ* before a catastrophic error occurs (see Methods for calculation of *δ*\*). Assuming that the estimation of each module becomes efficient before the joint estimation, Eq. 7 can be reinterpreted as a lower bound on the required decoding time before the estimation based on the joint likelihood function becomes efficient

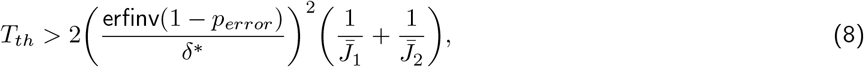

where erfinv(·) is the inverse of the error function and 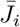 is the time-normalized Fisher information of module *i* (see Methods for derivation). Thus, the spatial periods of the modules influence the minimal decoding time by determining: (1) the largest allowed distance *δ*\* between the estimates of the modules, and (2) the variance of the estimations by the inverse of their respective Fisher information.

For example, if the spatial periods of the modules are very different, λ_2_ ≪ λ_1_, then there exist many peaks of *Q*_2_ around the correct peak of *Q*_1_ (Fig. 1c, top panel). More damaging still is the chance of having other modes of *Q*_1_ and *Q*_2_ close together. Thus, λ_2_ ≪ λ_1_ can create a highly multi-modal joint likelihood function (Wei et al. (2015) or Fig. 1c) where small deviations in 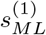 and 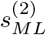 can cause a shift, or a change, of the maximal mode of the joint likelihood. To avoid this, |*δ*| < *δ*\* must be small leading to longer decoding times by Eq. 6. If, instead, the two modules have similar spatial periods λ_2_ ~ λ_1_, or λ_1_ is close to a multiple of λ_2_, then the distance between the peaks some period away are also close together (Fig. 1d) again leading to small *δ*\* and longer decoding time. Thus, assuming λ_1_ < 1, both small and large scale factors *c* can lead to long decoding times. In other words, periodic tuning suffers from the dilemma that small shifts in the individual stimulus estimates can cause catastrophic shifts in the joint likelihood function. For single-peaked tuning curves (λ_1_ = 1), however, only small scale factors *c* can pose such problems.

To test the approximation, we simulated a set of populations (*N* = 600 neurons) with different spatial periods. The populations were created using identical tuning parameters except for the spatial periods, whose distribution varied across the populations, and the amplitudes, which were adjusted to ensure an equal average firing rate (across all stimulus conditions) for all neurons (see Method for details on simulations and SI for the amplitudes). As described above, the spatial periods were related by a scaling factor *c*. Different values of *c* were tested for the largest period being either λ_1_ = 1 or λ_1_ = 1/2. Furthermore, only populations with unambiguous codes over the stimulus interval were included (Mathis et al., 2012). The minimal decoding time was found for each population by gradually increasing the decoding time until the empirical MSE was within two times the predicted lower bound (see Methods for details). Limiting the probability of catastrophic errors to *p_error_* = 10^−3^, Eq. 8 was found to be a good predictor of of the minimal decoding time (Fig. 1e, *R*^2^ = 0.89 and *R*^2^ = 0.95 for λ_1_ = 1 and λ_1_ = 1/2, respectively). In the case of λ_1_ = 1, the minimal decoding time is monotonically increasing with decreasing spatial periods (scale factor *c*) of module 2 (Fig. 1e, blue). For λ_1_ = 1/2, minimal decoding time is higher than that required for λ_1_ = 1. For λ_1_ = 1/2, the minimal decoding time increase with decreasing spatial periods. However, this trend is interrupted by large peaks (Fig. 1e, red line). The irregular trend is explained by the irregular behavior of *δ*\* (Fig. 1e top-left inset), whenever λ_1_ = 1/2 is close to a multiple of λ_2_, the smallest allowed deviation *δ*\* is also determined by the distance between peaks far away from the stimulus condition (Fig. 1c-d). Thus, whenever *c* is close to 1, 1/2, 1/3, etc., small displacements of the peaks of *Q*_1_ and *Q*_2_ can cause catastrophic shifts in the estimation of *s*_0_. Fig. 1 suggests that single-peaked tuning curves could remove catastrophic errors faster than any multi-peaked population.

### Minimal decoding times for more than two module populations

From the two-module case above, it is clear that the minimal decoding time becomes large whenever *c* << 1 or additionally, in the case of λ_1_ < 1, when the scale factor *c* is approximately 1, 1/2, 1/3, etc. However, Eq. 8 is difficult to analyse w.r.t. the scale factor *c* and is only valid for two module systems (*L* = 2). To approximate how the minimal decoding time scales with the distribution of spatial periods in populations with more than two modules, we extended the approximation method first introduced by Xie (2002). The method was originally used to assess the number of neurons required to reach the Cramér-Rao bound for single-peaked tuning curves with additive Gaussian noise for the ML-estimator. In addition, it only considered encoding a one-dimensional stimulus variable. We adapted this method to approximate the required decoding time for stimuli with an arbitrary number of dimensions, Poisson distributed spike counts, and tuning curves with arbitrary spatial periods. In this setting, the minimum decoding time can be approximated as (see Methods for derivation):

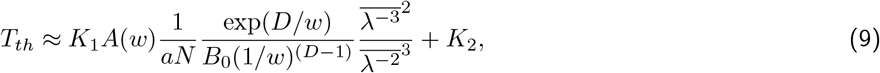

where the bars indicate the average w.r.t. the spatial periods in the population, *A*(*w*) is a function of *w* (see Method for detailed expression), and *K*_1_ and *K*_2_ are some unknown constants. Thus, Eq. 9 approximates how the minimal decoding time depends on the distribution of spatial periods. The derivation was carried out assuming the absence of spontaneous activity and that the amplitudes within each population are similar, *a*_1_ ≈ … ≈ *a_N_*. Importantly, the approximation assumes the existence of a unique solution to the maximum likelihood equations. It is therefore ill-equipped to predict the issues of stimulus ambiguity. Thus, it cannot capture the additional effects of λ_2_ ≪ λ_1_ or when λ_1_ is close to a multiple of λ_2_, as in Fig. 1c-d. On the other hand, complementing the theory presented in Eq. 8, Eq. 9 provides a more interpretable expression of the scaling of minimal decoding time. Assuming an equal number of neurons per module (and thus per spatial period), we can rewrite Eq. 9 in terms of *c* as

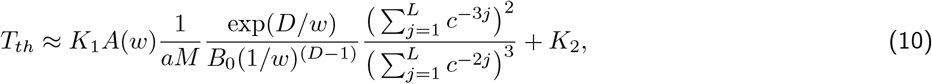

where *M* is the number of neurons in each module. Thus, for 0 < *c* ≤ 1, the minimal decoding time, *T_th_*, is expected to increase roughly linearly with decreasing scale factor, *c*. It also suggests that the scaling of the minimal decoding time with the scale factor should be similar for different choices of λ_1_. Furthermore, keeping the amplitudes fixed and increasing the stimulus dimensionality, *D*, should dramatically increase the minimal decoding time. Assuming all other parameters except *D* are constant, the minimal decoding time should grow roughly exponentially with the number of stimulus dimensions (see Eq. 9).

To confirm the validity of Eq. 10 we simulated populations of *N* = 600 tuning curves across *L* = 5 modules, where the spatial periods were again defined using a scale factor *c* and the largest period, λ_1_ (Fig. 2a). To avoid the effects of *c* ≪ 1, we limited the range of the scale factor to 0.3 ≤ *c* ≤ 1. The upper bound on *c* was kept to include entirely single-peaked populations, even though Eq. 10 might be a poor approximation for λ_1_ < 1 and *c* ≈ 1. Again, the assumption of homogeneous amplitudes in Eq. 10 was dropped in simulations (Fig. 2b, left column) to ensure that the average firing rate across the stimulus domain is equal for all neurons (Fig. 2b, right column). This had little effect on Fisher information, where the theoretical prediction was based on the average amplitudes across all populations with the same λ_1_ and stimulus dimensionality *D* (Fig. 2c inset, coloured lines are estimated Fisher information based on simulations and dashed black lines are the theoretical predictions using homogeneous amplitudes). As before, Fisher information grows with decreasing scale factor *c* and with decreasing spatial period λ_1_ (Fig. 2c, inset). However, increasing the stimulus dimensionality decreases Fisher information if all other parameters are kept constant. On the other hand, the minimal decoding time also increases with decreasing spatial periods (Fig. 2c). Furthermore, increasing stimulus dimensionality also increased the minimal decoding time. Using Eq. 10, the constants *K*_1_ and *K*_2_ were fitted using least square regression across populations sharing the same largest period, λ_1_, and stimulus dimensionality, *D*. Within this range of scale factors, Eq. 10 provides reasonable fits for the populations with λ_1_ = 1 (Fig. 2c, *R*^2^ ≈ 0.89 and *R*^2^ ≈ 0.90 for *D* = 1 and *D* = 2, respectively). For the populations with λ_1_ = 1/2, Eq. 10 becomes increasingly unable to predict the behavior of the minimal decoding time as *c* approaches 1 (see the red and yellow lines). On the other hand, as was suggested above, the scaling of the minimal decoding time with *c* is in fact similar for λ_1_ = 1 and λ_1_ = 1/2 whenever the scaling factor is less than ≈ 0.9 (Fig. 2c, compare the blue / red lines or the green / yellow lines). As suggested by Fig. 2d, there is also a strong correlation between Fisher information and minimal decoding time again indicating a speed-accuracy trade-off. As we will argue in the Discussion, the correlation between minimal decoding time and Fisher information is not simply due to a tougher requirement on MSE (from the Cramér-Rao bound) but reflects an important trade-off between accuracy and speed.

**Figure 2:**
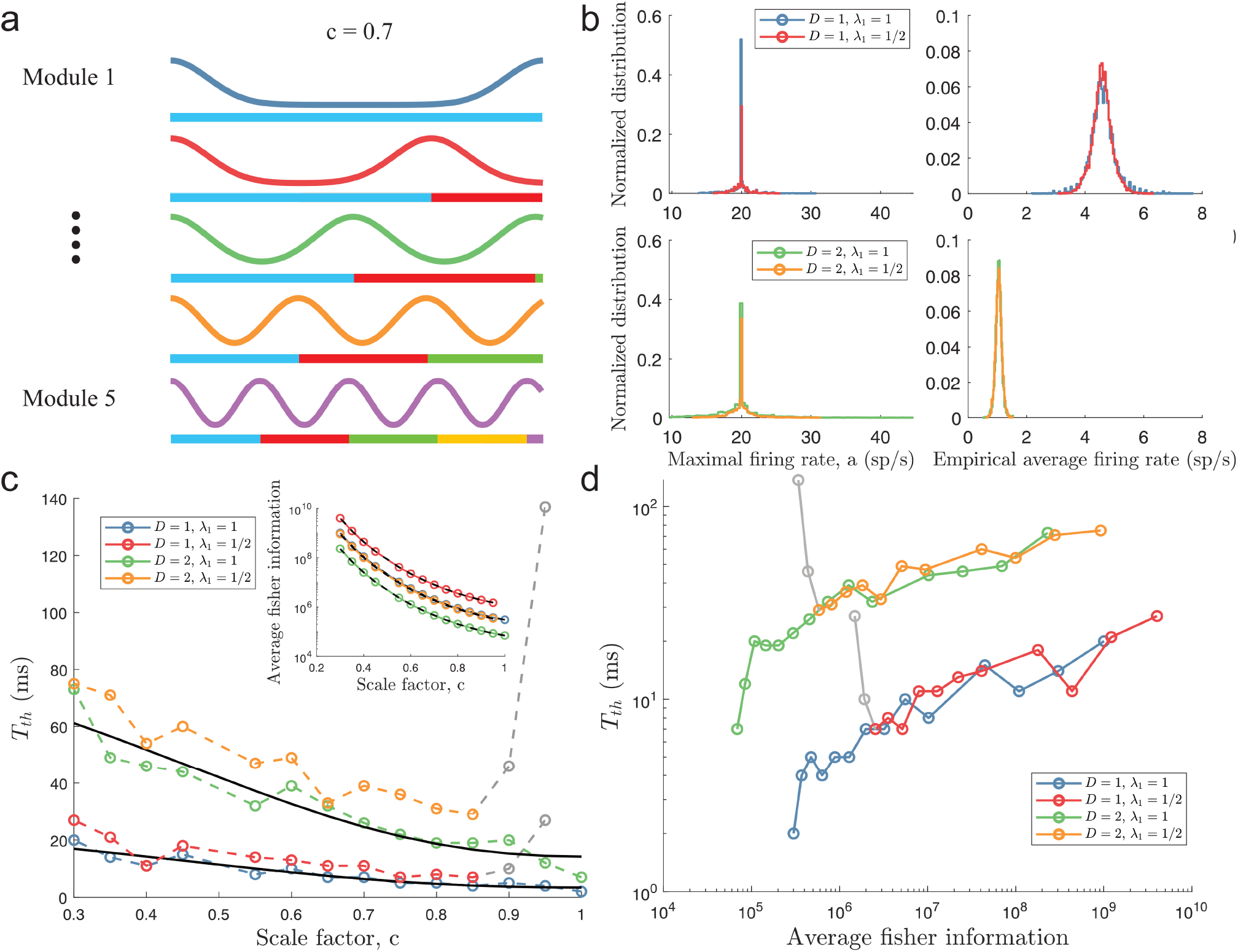
Minimal decoding times beyond two module populations. a) Illustration of the likelihood functions (for a single trial) of a population with *L* = 5 modules, where the spatial scale of each module relates to the previous module’s scale using a scale factor *c*. The code provides precise information about the stimulus (given sufficient decoding time) if *c* ≠ 1,1/2,1/3, etc. b) The peak amplitudes of each neuron (left column) were selected such that all neurons shared the same expected firing rate for a given stimulus condition (right column). Note that the reduction in average firing rates (across the stimulus domain) for *D* = 2 is not due to a reduction in peak amplitudes. c) Inset: Plot of average Fisher information as a function of the scale factor *c*. As before, smaller spatial scales imply larger Fisher information. Increasing stimulus dimensionality decreases information. The colored lines correspond to the estimated Fisher information in simulations, and the black dashed lines the theoretical predictions. Main plot: Plot of minimal decoding time as a function of scale factor *c*. Minimal decoding time tends to increase with decreasing grid scales (colored lines show the estimated minimal decoding time from simulations and the black lines show the fitted theoretical predictions). The gray color corresponds to points with large discrepancies between the predicted and the simulated minimal decoding times, likely a consequence of *c* being close to 1 (see Fig. 1c). d) Plot of the average Fisher information against the minimal decoding time. Shows a correlation between high Fisher information and long minimal decoding times (note the log-log scale). Points colored in gray are the same as in panel c). A list of all tuning parameters is given in Table S2.

Thus, while periodic tuning curves provide lower estimation errors for long decoding times by minimizing local errors (Fig. 2c, inset), a population of single-peaked tuning curves is faster at producing a statistically reliable signal by removing catastrophic errors (see Eq. 9 and Fig. 2c). Generalizing minimal decoding times to an arbitrary number of stimulus dimensions reveals that the minimal decoding time also depends on the stimulus dimensionality (see Fig. 2c, compare lines for *D* = 1 and *D* = 2). Interestingly, however, the approximation predicts that although minimal decoding time grows with increasing stimulus dimensionality, the minimal required spike count might be independent of stimulus dimensionality, at least for populations with integer spatial frequencies, i.e., integer number of peaks (see SI). The populations simulated here have non-integer spatial frequencies, but the trend of changes in mean spike count is still just slightly below 1 (indicating that slightly fewer spikes across the population were needed with increasing *D*, see Fig. S1). Thus, as the average firing rate decreases with the number of encoded features *D* (Fig. 2b), the increase in minimal decoding time with *D* can be largely explained by requiring a longer time to accumulate the sufficient number of spikes across the population.

### Effect of spontaneous activity

Many cortical areas exhibit spontaneous activity, i.e., activity that is not stimulus-specific (Snodderly and Gur, 1995; Barth and Poulet, 2012). Thus, it is important to understand the impact of spontaneous activity on minimal decoding time, too. Unfortunately, because our approximation of minimal decoding times did not include spontaneous activity, we relied on simulations to study the effect of such non-specific activity.

When including independent ongoing spontaneous activity at 2 spikes/second to all neurons for the same populations as above, minimal decoding times were elevated across all populations (Fig. 3). Furthermore, the minimal decoding time increased faster with decreasing *c* in the presence of spontaneous activity compared to the case without spontaneous activity (ratios of fitted *K*_1_ in Eq. 10 were approximately 1.69 and 1.72 for *D* = 1 and *D* = 2, respectively). Thus, spontaneous activity can have a substantial impact on the time required to produce reliable signals. Fig. 3 also suggests that areas with spontaneous activity are less suited for periodic tuning curves. Especially, the combination of multidimensional stimuli and spontaneous activity leads to much longer minimal decoding times for tuning curves with small spatial periods (*c* < 1 or λ_1_ < 1). For example, when encoding a two-dimensional stimulus, only the populations with (λ_1_ = 1, *c* =1) and (λ_1_ = 1, *c* = 0.95) could remove catastrophic errors in less than 40 ms when spontaneous activity at 2 spikes/second was present. Thus, the ability to produce reliable signals at high speeds severely deteriorates for periodic tuning curves in the presence of non-specific spontaneous activity.

**Figure 3:**
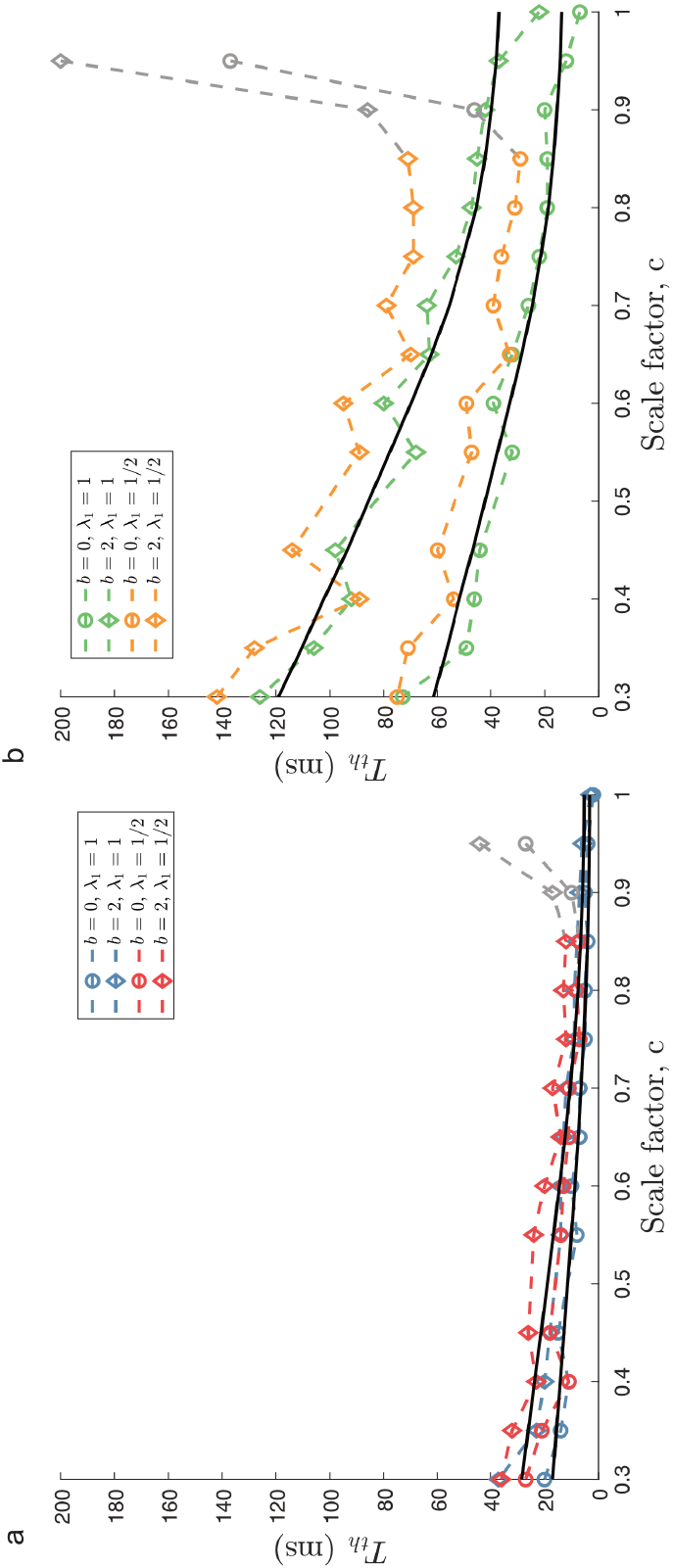
Effect of spontaneous activity. a) The case of encoding a one-dimensional stimulus (*D* = 1) with or without spontaneous activity at 2 spikes/second (diamond and circles shapes, respectively). b) The case of a two-dimensional stimulus (*D* = 2) under the same conditions as for a). Spontaneous activity increases the time required for all populations to produce reliable signals, but the effect is strongest for *c* ≪ 1. All other tuning curve parameters are set according to Table S2.

This result has an intuitive explanation. The amount of catastrophic errors depends on the probability that the trial variability reshapes the neural activity to resemble the possible activities for a distinct stimulus condition (see Fig. 1a). From the analysis presented above, periodic tuning curves have been suggested to be more susceptible to such errors. Adding spontaneous activity does not reshape the tuning curves themselves but only increases the trial-by-trial variability. Thus, by this reasoning, it is not surprising that the systems which already were more susceptible suffered more strongly from the increased variability induced by spontaneous activity. The importance of Fig. 3 is that even spontaneous activity as low as 2 sp/s can have a clearly visible effect on minimal decoding time.

## Implications for a simple spiking neural network with sub-optimal readout

To further illustrate the relationship between the shape of tuning curves and minimum decoding time, we simulated simple two-layer feed-forward spiking neural networks to decode time-varying stimulus signals. The first layer corresponds to the tuning curves, and the neurons were kept unconnected. The stimulus-specific tuning of the inputs to these neurons is either fully single-peaked, creating a population of single-peaked tuning curves, or periodic with different spatial periods, creating a population of periodic tuning curves (Fig. 4a). Given their tuning preferences, neurons in the first layer responded with rate modulated Poisson type spike trains. The modulation strengths of the inputs were chosen to ensure that the average input to each neuron was equal. The second layer instead acted as a readout layer. This layer received both stimulus-specific excitatory input from the first layer and external non-specific excitation (corresponding to background activity). The connection strength between the first and second layers depended on the difference in preferred stimulus conditions between the pre- and post-synaptic neurons. Such connectivity could, for example, be obtained by unsupervised Hebbian learning. Because the tuning curves in the first layer can be periodic, they can connect strongly to several readout neurons. We introduced lateral inhibition among the readout neurons (without explicitly modeling inhibitory neurons) to create a winner-take-all style of dynamics, where the readout neurons with large differences in preferred stimulus inhibit each other more strongly. Decoding is assumed to be instantaneous and based on the preferred stimulus condition of the spiking neuron in the readout layer. We tested two different types of time-varying stimuli: (1) a step-like change from *s* = 0.25 to *s* = 0.75 (Fig. 4b, blue trace) and (2) a continuously time-varying stimulus drawn from an Ornstein–Uhlenbeck process (Fig. 4c blue trace; see Methods). The stimulus was instantaneously decoded whenever a readout neuron spikes, and the estimate was compared to the current true stimulus value (see Fig.4b-c for individual trials).

**Figure 4:**
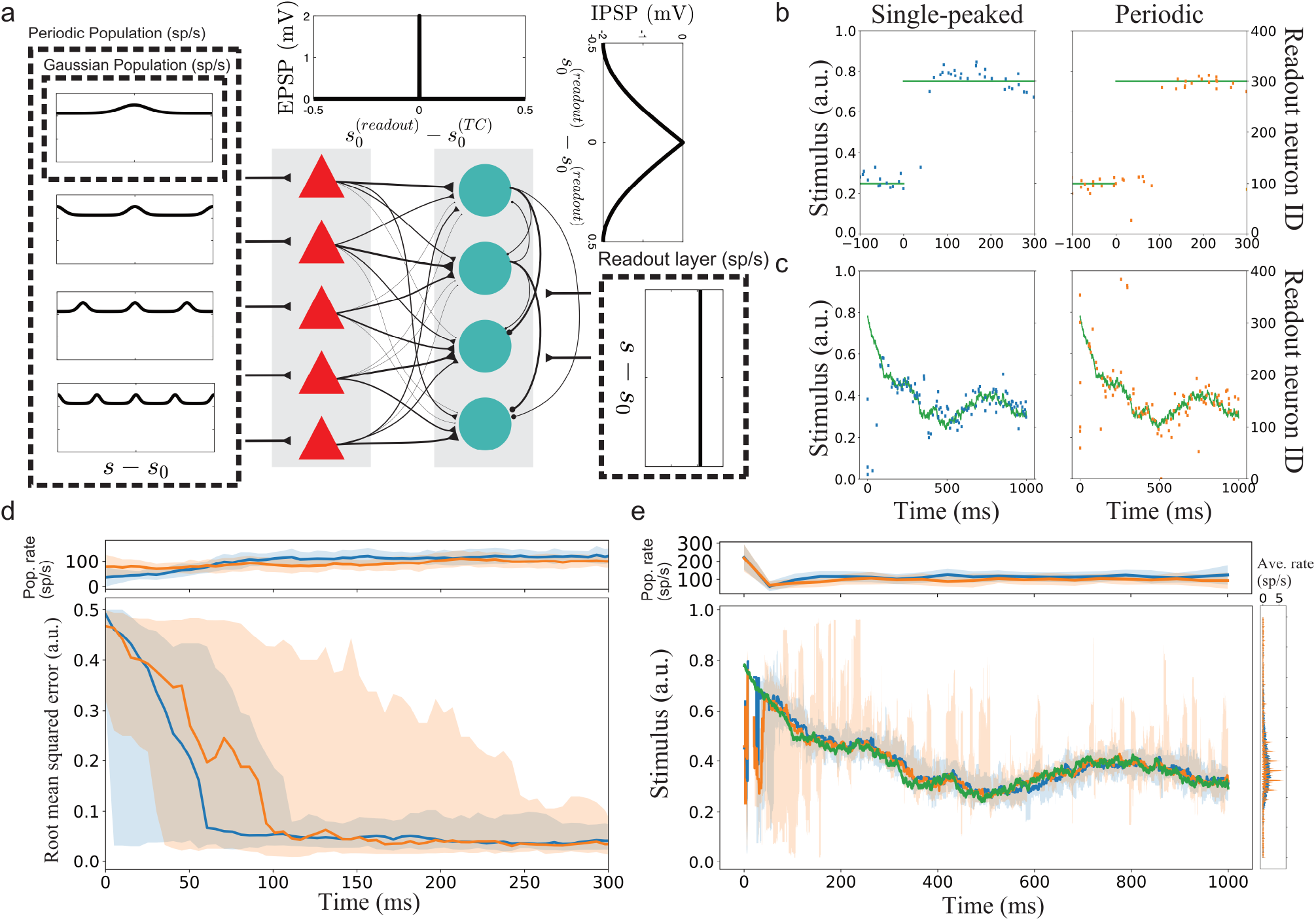
a) Illustration of the spiking neural networks (SNNs). b) Two example trials for step-like change in stimulus (blue line). The left and right plots show the readout activity (red) for the single-peaked and periodic SSNs, respectively. Note that the variance around true stimulus is larger for the single-peaked SNN (i.e., larger local errors) but that there are fewer catastrophic errors than for the periodic SNN. c) Same as for b) but with a continuously time-varying stimulus. d) Main panel: The median RMSE (thick lines) over all trials in a sliding window (length 50 ms) for the single-peaked (blue) and periodic (orange) SNNs. The shadings correspond to the regions between the 5th and 95th percentiles. Top panel: The instantaneous population firing rates of the readout layers and the standard deviations (the color code is same as in main panel). e) The median estimated stimulus over all trials in a sliding window (length 10 ms) for the single-peaked (blue) and periodic (orange) SNNs. Shaded areas again correspond to the regions between the 5th and 95th percentiles. The true stimulus is shown in green.

In the case of a step-like stimulus change, the readout layer for the single-peaked population required a shorter time to switch states than the periodic network (see Fig. 4d). The shorter switching time is consistent with the hypothesis that single-peaked tuning curves have shorter minimal decoding times than periodic tuning curves. In these simulations, the difference is mainly due to some neurons in the first layer of the periodic network responding both before and after the step change. Thus, the correct readout neurons (after the change) must compensate for the hyper-polarization built up before the change and the continuing inhibitory input from the previously correct readout neurons (which still get excitatory inputs, too). Note that there are only minor differences in the population firing rates between the readout layers, suggesting that this is not a consequence of different excitation levels but rather of the structures of excitation.

The continuously time-varying stimulus could be tracked fairly well by the network with single-peak or periodic tuning curves. However, averaging across trials showed that SNNs with periodic tuning curves have larger sporadic fluctuations (Fig. 4e). This suggests that decoding with periodic tuning curves has difficulties in accurately estimating the stimulus without causing sudden, brief periods of large errors. To make a statistical comparison between the populations, we investigated the distributions of root mean squared error (RMSE) across trials. In both stimulus cases, there is a clear difference between the network with single-peaked tuning curves and the network with periodic. For the step-like change in stimulus condition, a significant difference in RMSE arise roughly 100 ms after the stimulus change (Fig. 5a, using two-sample Kolmogorov–Smirnov (KS) test based on 30 trials per network). For the time-varying stimulus, using single-peaked tuning curves also results in significantly lower RMSE compared to a population of periodic tuning curves (Fig. 5b, p-value < 0.001 using a two-sample KS test based on 30 trials per network, RMSE calculated across the entire trial).

**Figure 5:**
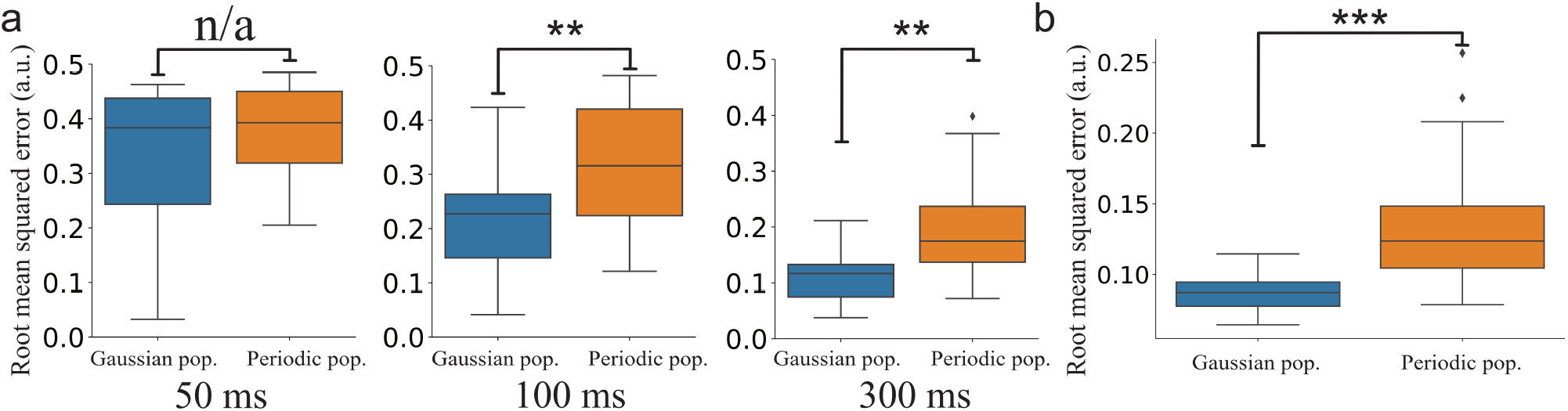
a) Statistical comparison between the distributions of cumulative RMSEs at different decoding times (p-values 0.4, 9.0 · 10^−4^, and 8.7· 10^−5^, respectively). b) The distributions of RMSE across trials for the two SNNs (p-value = 4.3 · 10^−8^).

## Discussion

Several studies have suggested that periodic tuning creates an unparalleled precise neural code by minimizing local errors (Sreenivasan and Fiete, 2011; Mathis et al., 2012; Wei et al., 2015). Nevertheless, despite this advantages of periodic tuning, single-peaked tuning curves are widespread in early sensory areas especially in the early visual system. Is the single-peaked tuning simply sub-optimal, or are there other factors that can favor the ubiquity of single-peaked tuning in early sensory processing? Despite a long history of studying information representation using rate-based tuning curves, the effect of spatial periodicity and catastrophic errors on the required decoding time has not been addressed. Here, we show that the possibility of catastrophic estimation errors (Fig. 1a) introduces the possibility that different shapes of tuning curves can have different minimal decoding times.

But why study the minimal decoding time and not simply compare the time-evolution of the MSE directly instead? However, comparing MSE directly between populations can be a misleading measure of reliability if the distributions of errors are qualitatively different. That is, if the amounts of local errors are different, lower amounts of catastrophic errors do not necessarily imply lower MSE (see Fig. S2 for an example). Thus, a comparison of MSE only becomes valid once the minimal decoding times have been met. Here we assume that catastrophic errors should strongly affect the usability of a neural code. Therefore, we argue that the first criterion for any rate-based neural code should be to satisfy its constraint on decoding time. This led to the following question: is there a trade-off between the accuracy (i.e., Fisher information) of a neural code and the minimal required decoding time for single-peaked and periodic tuning?

The answer is yes. We found that minimal decoding time increased with with decreasing the spatial periods of tuning curves (Fig. 2c), suggesting a trade-off between accuracy and speed for populations of tuning curves. Experimental data suggest that minimal decoding times can be very short, of the order of tens of milliseconds, reflecting that a considerable part of the information contained in firing rates over long periods is present in short sample periods, too (Tovee et al., 1993). Moreover, the first few spikes have been shown to carry significant amounts of task information in both visual (Resulaj et al., 2018) and olfactory areas (Resulaj and Rinberg, 2015). In our simulations, tens of spikes carry enough information to produce a reliable estimate of the stimulus free of catastrophic errors. As with decoding time, single-peaked tuning curves also need fewer spikes to produce reliable signals. Thus, the speed-accuracy trade-off can be reinterpreted as a trade-off between being accurate or efficient. In simulated networks with spiking neurons, we showed that the use of periodic tuning curves increased the chances of large instantaneous estimation errors, leading to longer times before switching “states” (Fig. 4d) and difficulties tracking a time-varying stimulus (Fig. 4e).

The notion of speed-accuracy trade-off is further strengthened for high-dimensional stimuli in the sense that, when encoding a high-dimensional stimulus, using the same distribution of spatial periods leads to an increase in minimal decoding time. Natural stimuli have higher dimensions than typically used in animal experiments. Many sensory neurons are tuned to multiple features of the external stimulus, creating such mixed selectivity of features (e.g., Garg et al. (2019)). For neurons responding to task-related variables, mixed selectivity has been shown to enable linear separability and to improve discriminability (Rigotti et al., 2013; Fusi et al., 2016; Jeffrey Johnston et al., 2020). For continuous stimulus estimations, mixed selectivity has also been proposed to decrease MSE when decoding time is limited (Finkelstein et al., 2018). However, to remove catastrophic errors, which as we have argues is not necessarily synonymous with lower MSE, the exponential increase in minimal decoding time could easily lead to very long decoding times, on the order of seconds, even for stimulus with dimensionality not higher than 4 or 5. Thus, minimal decoding time should set a bound on the number of features a population can jointly encode reliably. In addition, neurons in sensory areas often exhibit a degree of non-specific activity (Snodderly and Gur, 1995; Barth and Poulet, 2012). Introducing spontaneous activity to the populations in our simulations further amplified the differences in minimal decoding times (Fig. 3). Thus, for jointly encoded stimuli, especially in areas with high degrees of spontaneous activity, a population of single-peaked tuning curves might be the optimal encoding strategy for rapid and reliable communication.

To conclude, we provide normative arguments for the single-peaked tuning of early visual areas. Rapid decoding of stimulus is crucial for the survival of the animals. Consistent with this, animals and humans can process sensory information at impressive speeds. For example, the human brain can generate differentiating event-related potentials to go/no-go categorization tasks using novel complex visual stimuli in as little as 150 milliseconds (Thorpe et al., 1996). These “decoding” times do not decrease for highly familiar objects, suggesting that visual processing is a highly automatized feed-forward process operating at speeds that cannot be reduced (Fabre-Thorpe et al., 2001). Given constraints on low latency communication, it is crucial that each population can produce a reliable signal fast. In this regard, single-peaked tuning curves are indeed superior compared to periodic. Thus, if the available decoding time is short, it is preferable to lower the accuracy over long sample periods in exchange for a signal which can be reliably produced within the desired time window. The fact that early visual areas exhibit spontaneous activity and encode multi-dimensional stimuli further strengthens the relevance of the differences in minimal decoding times. We note that these results might extend beyond the visual areas, too. For example, hippocampal place cells involved in spatial navigation (O’Keefe and Dostrovsky, 1971; Wilson and McNaughton, 1993) are known for their single-peaked tuning (but see Eliav et al. (2021)). The interesting observation in this context is that place cells produce more reliable signals than their input signals from the medial entorhinal cortex with a combination of single- and multi-peaked tuning (Cholvin et al., 2021).

In general, our work highlights that minimum decoding time is an important attribute of a neural code and should be considered while evaluating candidate neural codes. Our analysis suggests that decoding of high dimensional stimuli can be prohibitively slow with rate-based tuning curves. Experimental data on the representation of high-dimensional stimuli is rather scant as relatively low-dimensional stimuli are typically used in experiments (e.g., oriented bars). Our work gives a compelling reasons to understand whether and how biological brains can reliably encode high-dimensional stimuli at behaviorally relevant time scales.

## Methods

### Minimal decoding times - Simulation protocols

To study the dependence of decoding time *T* on MSE for populations with different distribution of spatial frequencies, we simulated populations of synthetic tuning curves (Eq. S.1). The stimulus was chosen to be circular with range [0, 1)^*D*^ to avoid boundary effects and the parameters of the tuning curves are given in Table S1-S2. The preferred stimulus conditions *s′* were sampled independently from a random uniform distribution over [0,1) (independently and uniformly for each stimulus dimension). The preferred locations *s′* were shared across all populations to ensure equal comparison. In each trial, a stimulus *s* ∈ [0,1)^*D*^ was also independently sampled from a uniform distribution over [0,1)^*D*^. The spike counts for each neuron were then sampled according to Eq.S.3.

Minimal decoding time was defined as the minimal decoding time for which the neural population approximately reaches the Cramér-Rao bound. To estimate the reaction time in simulations, we incrementally increased decoding time *T* (using 1 ms increments, starting at *T* = 1 ms) until

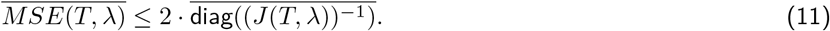

Note that the mean here refers to the mean across stimulus dimensions (for multi-dimensional stimuli) and that diag(·) refers to taking the diagonal elements from the inverse of the Fisher information matrix, (*J*(*T*, λ))^−1^. For a given decoding time *T*, the estimation of MSE was done by continually sample random stimulus conditions (from a uniform distribution), sample a noisy response to the stimulus (Poisson distributed spike counts) and then apply maximum likelihood estimation (see section ‘Implementation of maximum likelihood estimator’ for details on implementation). This was repeated until the first two non-zero digits of the MSE had been stable for 1000 consecutive random stimulus samples (see Alg. S1 in SI). Because the Fisher information matrix *J* was estimated only in the special case without spontaneous activity, it was in simulations approximated by the element-wise average across 10000 randomly sampled stimulus conditions (sampled according to the uniform distribution over the stimulus domain), where each element was calculated according to Eq. S.18 or Eq. S.19 given a random stimulus trial. A high-level view of the simulation of minimal decoding times is given in Alg. S2 (SI).

#### Implementation of maximum likelihood estimator

Given some noisy neural responses, **r**, the maximum likelihood estimator (MLE) chooses the stimulus condition which maximizes the likelihood function, 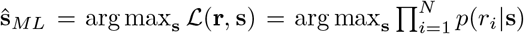. A common approach is to instead search for the maximum of the log-likelihood function (the logarithm is a monotonic function and therefore preserves any maxima/minima). The stimulus-dependent terms of the log-likelihood can then be expressed as

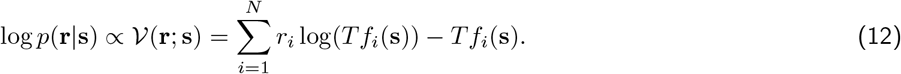

Unfortunately, the log-likelihood function is not guaranteed to be concave, and finding the stimulus condition 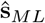 which maximizes the log-likelihood function is not trivial (a non-convex optimization problem). To overcome this difficulty, we combined grid-search with the Nelder–Mead method, an unconstrained non-linear program solver (implemented using MATLAB’s built-in function *fminsearch*, https://www.mathworks.com/help/matlab/ref/fminsearch.html). Grid search was used to find a small set of starting points with the largest log-likelihood values (in simulations, the four stimulus conditions with the largest log-likelihood values were used). The true stimulus condition **s**\* was always added into the set of starting points regardless of the log-likelihood value of that condition (yielding a total of 5 starting points). Then the Nelder–Mead method was used with these starting points to find a set of (possibly local) maxima. Thus, this approach does not overestimate the amount of threshold distortion but can potentially miss some global estimation errors instead. Given a estimated stimulus 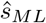, the error was then evaluated along each stimulus dimension independently

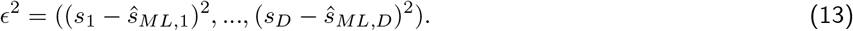

### Spiking network model

#### Stimuli

As in the previous simulations, we assumed that the stimulus domain was a circular stimulus defined between [0, 1). We simulated the responses to two different types of stimuli, (1) a step-like change in stimulus condition from *s* = 0.25 to *s* = 0.75 and (2) a stimulus drawn from a modified Ornstein–Uhlenbeck process

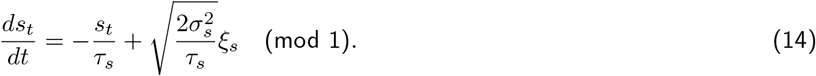

For parameter values, see Table S4.

#### Network model

The spiking networks were implemented as two-layer, feed-forward networks using LIF neurons (see Tab. S5 for parameter values). The neurons in the first layer were constructed to correspond to either single-peaked or periodic tuning curves. Two networks were tested, one network where the first layer corresponds to single-peaked tuning curves and a second network corresponding to periodic tuning curves (with *L* = 4 modules). For each neuron *i* in module *j* in the first layer, the input to was drawn from independent Poisson point processes with stimulus dependent rates 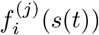

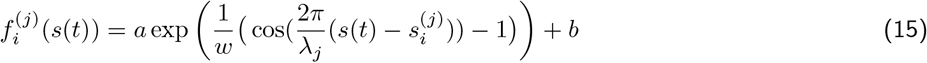

Here, the constants *a* and *b* were chosen such that the baseline firing rate was slightly above zero and the maximal firing rate was slightly below 20 sp/s (see Tab. S6 for all network related parameter values). Each pre-synaptic spike caused an EPSP of size *J_E_* in the first layer. For each module in the first layer, the preferred locations 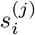 were equidistantly placed across [0, λ*_j_*). The neurons in the second layer were only tuned to a single preferred stimulus location each, equidistantly placed across [0, 1). Whenever a spike occurred in the first layer, it elicited EPSPs with a delay of 1.5 ms in all neurons in the second layer. The size of the EPSPs depended on the difference in preferred tuning (Δ) between the pre- and post-synaptic neurons

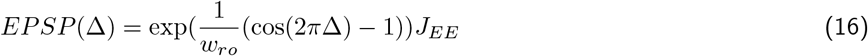

Here *J_EE_* determines the maximal EPSP (mV) and the constant *w_ro_* was chosen such that the full width at half maximum of the EPSP kernels tiled the stimulus domain without overlap. Note that for periodically tuned neurons (i.e., with multiple preferred locations) in the first layer, the smallest difference in preferred tuning was chosen for each neuron in the second layer.

As for the excitatory neurons in the first layer, whenever a spike occurred in the second layer, it elicited IPSPs with a delay of 1.5 ms in all other neurons in the second layer. Again, the size of the IPSPs depended on the difference in preferred tuning (Δ) between the two neurons, but this time according to

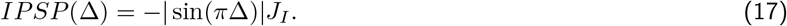

Thus, the range of inhibition was much broader compared to the excitation.

#### Evaluating decoding performance

We assumed that the decoder was instantaneously based on the neuron index of the firing neuron in the readout layer. Given that Φ(*t_k_*) is a function that provides the index of the neuron firing at time *t_k_*, the stimulus is instantaneously decoded to

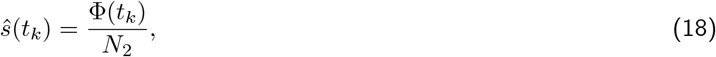

where *N*_2_ is the number of neurons in the readout layer. For both stimulus cases, the decoding performance was evaluated using (1) the distribution of RMSE (Fig. 4d) or estimated stimulus conditions (Fig. 4e) in a sliding window or (2) the distributions of accumulated RMSE (Fig. 5).

### Simulation tools

All the simulation were done using code written in MATLAB and Python (using Brian2 simulator (Stimberg et al., 2019)). The simulation code will be made available on Github upon publication of the manuscript.

### Approximating minimal decoding time in 2-module systems

To gain understanding of the interaction of two modules with different spatial periods, consider the likelihood function as a product of the likelihood functions of the two modules individually

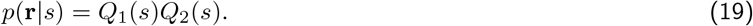

Using the Laplace approximation, each of these functions can be approximated as a periodic sum of Gaussians (Wei et al., 2015)

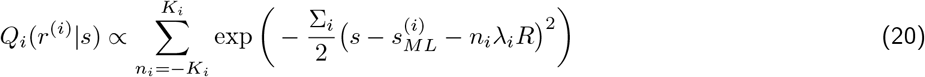

where 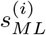 denotes the peak closes to the true stimulus condition *s*_0_ and 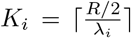 (note that for λ_1_ = 1, we set *K*_1_ = 0 to avoid repeating the same mode). Assuming that each module is efficient, the width of the Gaussians as can be approximated as

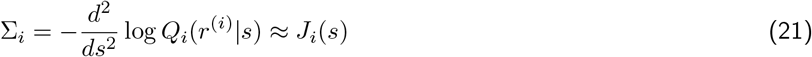

where *J_i_*(*s*) ≈ *J_i_* is the Fisher information of module *i*. The joint likelihood function can thus be approximated as

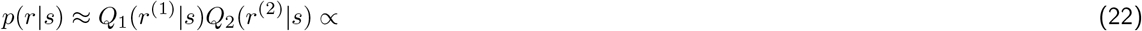

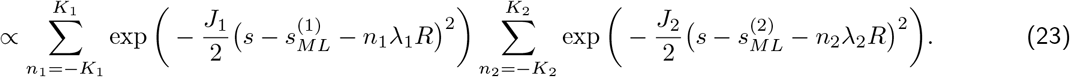

As the likelihood functions depend on the particular realization of the spike counts, the distance between the modes of the respective likelihoods closest to the true stimulus condition *s*_0_, 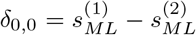, is a random variable. Note that in the result section, *δ*_0,0_ is simply referred to as *δ* for clarity.

The joint likelihood distribution *p*(*r*|*s*) has its maximal peak close to the true stimulus condition *s*_0_ if *δ*_0,0_ is the smallest distance between any pairs of peaks of *Q*_1_ and *Q*_2_. Assuming that both modules provide efficient estimates, the distance *δ*_0,0_ can be approximated as a normally distributed random variable

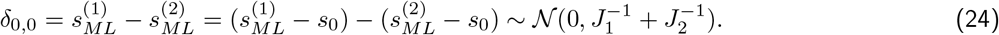

The distance between any pair of peaks in *Q*_1_ and *Q*_2_ within the stimulus range becomes

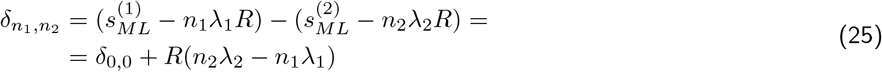

where *n*_1_ ∈ {–*K*_1_, …, *K*_1_} and *n*_2_ ∈ {–*K*_2_, …, *K*_2_} are indexing the different Gaussians as before. Thus, the threshold point for catastrophic error is if there is another pair of modes with same distance between them, i.e.,

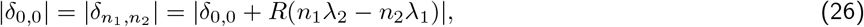

for some *n*_1_ and *n*_2_ belonging to the index sets as above. Thus, to avoid catastrophic errors, it is necessary that

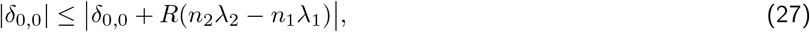

for all *n*_1_ ∈ {–*K*_1_, …, *K*_1_} and *n*_2_ ∈ {–*K*_2_, …, *K*_2_}. By solving Eq. 27, and taking into account that *R*(*n*_2_λ_2_ – *n*_1_λ_1_) can be either positive or negative, we get

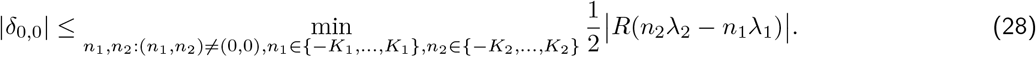

Assuming that the period of the second module is a scaling of the first module, λ_2_ = *c*λ_1_, the above equation becomes

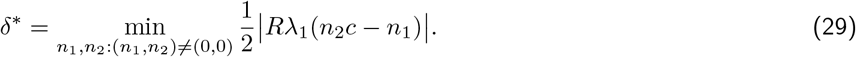

Note that stimulus ambiguity can never be resolved if *δ*_*n*_1_, *n*_2__ = *δ*_0,0_ for some pair (*n*_1_, *n*_2_) ≠ (0, 0), which is analogous to the condition in (Mathis et al., 2012).

To limit the probability of catastrophic estimation errors from the joint distribution to some small error probability *p_error_*, the following should hold

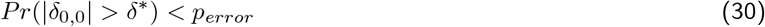

Because 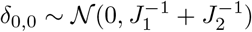, we have

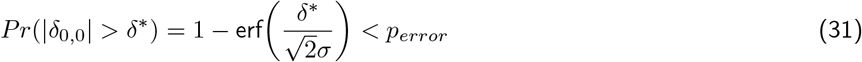

where erf(·) is the error-function and 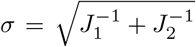. By rearranging the terms and using Eq.S.14, we can obtain a lower bound on the required decoding time

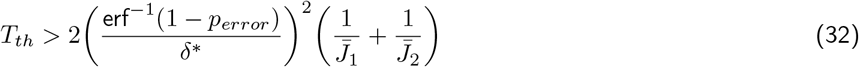

where 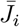 is the time-normalized Fisher information of module *i*. Note that *δ*\* can easily be found using an exhaustive search according to Eq. 28 or Eq. 29.

### Approximating minimal decoding time

To approximate the order by which the population reaction time scales with the distribution of spatial frequencies and the stimulus dimensionality, we extended the approximation method introduced by Xie (2002). The key part of the approximation method is to use a Taylor series to reason about which conditions must hold for the distribution of errors to be normally distributed with a covariance equal to the inverse of the Fisher information matrix. Note that this approximation assumes the existence of a unique solution to the maximum likelihood equations, thus it is not applicable to ambiguous neural codes (e.g., *c* = 1/2,1/3,1/4,… etc).

First, let’s recollect the Taylor series with Lagrangian reminder for a general function g

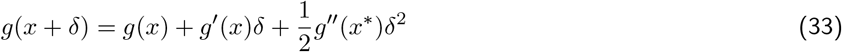

where *x*\* is somewhere on the interval [*x, x* + *δ*). Thus, in the multivariate case, the derivative in the j:th direction of the log-likelihood function for stimulus condition 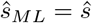 can be rewritten using a Taylor series with Lagrangian reminder as

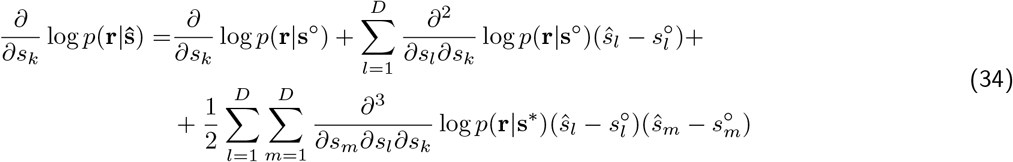

for all *k* ∈ {1, …, *D*} where **s**° is the true stimulus condition and **s**\* is a stimulus point between **s**° and 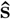.

If the estimated stimulus is close to the true stimulus then the quadratic order terms are small. If so, the variance of 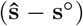 converges towards 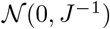 (in distribution), where *J* is the Fisher information matrix (Lehmann and Casella, 1998). However, if the estimated stimulus in not close to the true stimulus, then the quadratic terms are not negligible. Therefore, when *T* is sufficiently large and the variance of the estimation follows the Cramér-Rao bound, the following should hold for all *k* ∈ {1, …, *D*}

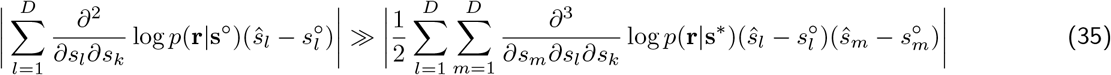

In this regime, we do the following term-wise approximations

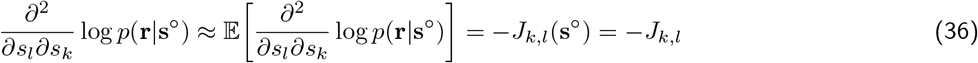

and

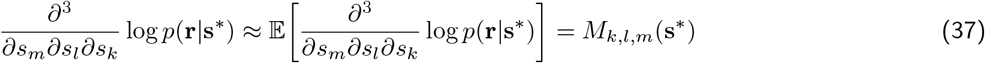

which gives

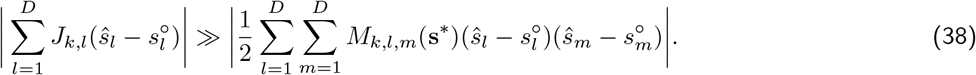

Because *M_k,l,m_* ≈ 0 unless *k* = *l* = *m* (see SI) and using an upper bound for 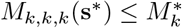 for all **s**\*, Eq. 38 simplifies to

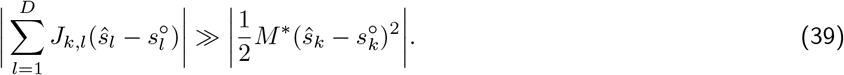

Furthermore, because *J*(**s**) is a diagonal matrix (see SI), we have

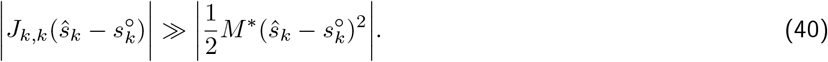

Next, by taking the square of the absolute values, we obtain

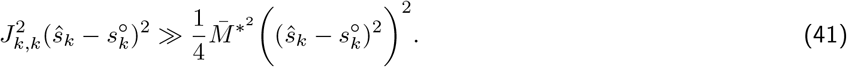

Because we assumed that *N* and *T* are sufficiently large to meet the Cramér-Rao bound, we have that

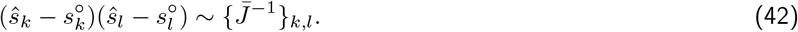

Inserting Eq. 42 into Eq. 41 gives

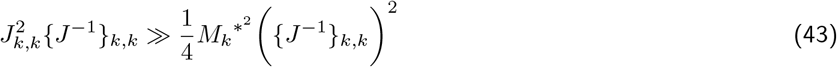

or, equivalently,

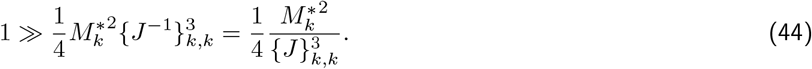

By approximating the term 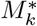 (see SI) and using the expression for Fisher information (Eq.S.14), the expression for population reaction times can be obtained as

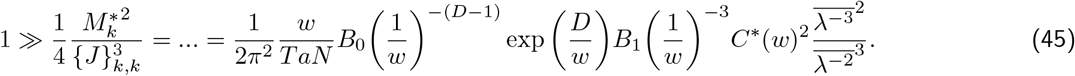

Where *C*\*(*w*) is a undetermined function of the width parameter *w*. By reorganizing the expression, one obtains

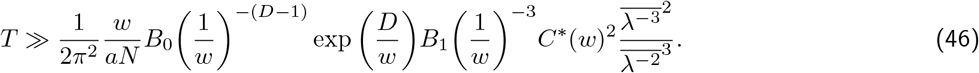

As the validity of the approximation decreases with *w* (see SI), we only collect the terms which includes *D*, *ξ*, *a* and *N*, and approximate the population reaction time as

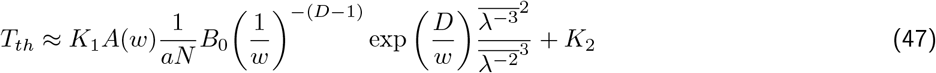

where *A*(*w*) is some unknown function of *w* and *K*_1_ and *K*_2_ are constants.

