## Supplemental Information for "How short decoding times, stimulus dimensionality and spontaneous activity constrain the shape of tuning curves: A speed-accuracy trade-off"

#### 1 Tuning curves and spike count model

In the paper, we study the representation of a multidimensional stimulus $s = (s_1, ..., s_D)$. For simplicity, it is assumed that the range of the stimulus in each dimension is equal, such that $s_j \in [0, R)$ for all $j \in \{1, ..., D\}$. Note that this assumption do not qualitatively change the results. Furthermore, we assume that the tuning curves were circular (von Mises) tuning curves

$$f_i(s) = a_i \prod_{j=1}^{D} \exp\left(\frac{1}{w}\big(\cos(\frac{2\pi}{\lambda_i R}(s_j - s'_{j,i})) - 1\big)\right) + b = a_i \prod_{j=1}^{D} q_{j,i}(s) + b, \tag{S.1}$$

where $a_i$ is the peak amplitude of the stimulus-related tuning curve of neuron $i$, $w$ is a width scaling parameter, $\lambda_i$ defines the spatial period of the tuning, $s_{j,i}$ determines the location of the firing field(s) in the $j$:th dimension, $D$ is the number of stimulus dimensions, and $b$ determines the amount of background activity. The amplitude parameters $a_i$ were tuned such that all tuning curves had the same firing rate when averaged across all stimulus conditions (see Eq. S.6).

It is possible to reparametrize the stimuli into a phase variable, $\phi = \frac{s_j}{R}$. In the article, calculations and numerical simulation are based on phase variables $\phi$. This only changes the MSE and fisher information by a constant scaling $\frac{1}{R^2}$. As we are interested in comparing the minimal decoding time, not the MSE, we can drop the "unnormalized" stimulus $s$. The tuning curves in Equation S.1 can thus be rewritten using the phase variable $\phi$ as

$$f_i(\phi) = a_i \prod_{j=1}^{D} \exp\left(\frac{1}{w}\big(\cos(\frac{2\pi}{\lambda_i}(\phi_j - \phi'_{j,i})) - 1\big)\right) + b = a_i \prod_{j=1}^{D} q_{j,i}(\phi) + b. \tag{S.2}$$

Given stimulus condition $s$ and decoding time $T$, the spike count of each neuron was independently sampled from a Poisson distribution with rate $Tf_i(s)$. Thus, the probability of observing a particular spike count pattern $\mathbf{r} = (r_1, ..., r_N)$ given $s$ is

$$p(\mathbf{r}|\mathbf{s}) = \prod_{i=1}^{N} p(r_i|\mathbf{s}) = \prod_{i=1}^{N} \frac{\big(Tf_i(\mathbf{s})\big)^{r_i} \exp\big(-Tf_i(\mathbf{s})\big)}{r_i!}. \tag{S.3}$$

##### 1.1 Adjusting amplitudes

In order to make a fair comparison of decoding times across populations, we constrain each neuron to have the same average spike count per unit time across the stimulus domain. The spike count for a neuron and a given stimulus condition is

$$\mu_i(s) = \mathbb{E}_{r_i}\left[r_i \middle| s\right] = Tf_i(s) \tag{S.4}$$

where $T$ is the finite time window. Thus, the average spike count (for $T = 1$ second) over both stimulus conditions and trials is

$$\mu_i = \mathbb{E}_s\left[\mathbb{E}_r\left[r_i \middle| s\right]\right] = \left\{\text{Stimulus Uniformly distributed}\right\} = b + a_i \int_0^1 q_i(\phi) d\phi. \tag{S.5}$$

Thus, given a desired stimulus-related spike count, $\mu^*$, the amplitudes will be set to

$$a_i = \frac{\mu^*}{\int_0^1 q_i(\phi)d\phi}. \tag{S.6}$$

Note that the integrals in Equation S.6 are analytically solvable whenever the relative spatial frequency $\xi_i = 1/\lambda_i$ is a positive integer, in which case we have

$$\prod_{j=1}^{D}\int_0^1 \exp\left(\frac{1}{w_i}\big(\cos(\frac{2\pi}{\lambda_i}(\phi_j-\phi'_{j,i}))-1\big)\right)d\phi = B_0\left(\frac{1}{w_i}\right)^D \exp\left(-\frac{D}{w_i}\right). \tag{S.7}$$

regardless of $\phi'_i$, here $B_0(\cdot)$ is the modified Bessel function of the first kind. In simulations, $\mu^*$ was set such that tuning curves with integer spatial frequencies $(1/\lambda)$ have amplitudes of $20$ sp/s.

### 2 Fisher information and the Cramér-Rao bound

Assuming a one-dimensional variable, the Cramér-Rao bound gives a lower bound on the MSE of any estimator $G$ (Cover and Thomas, 2005)

$$\mathbb{E}[(G(\mathbf{r})-s)^2] \geq \frac{[1+b'_G(s)]^2}{J(s)} + b_G(s)^2, \tag{S.8}$$

where $b_G(s) = \mathbb{E}[G(\mathbf{r})-s]$ is the bias of the estimator $G$ and $J(s)$ is fisher information, defined as

$$J(s) = \mathbb{E}\left[\frac{\partial}{\partial s}\log p(\mathbf{r}|s)\right]^2 = -\mathbb{E}\left[\frac{\partial^2}{\partial s^2}\log p(\mathbf{r}|s)\right] \tag{S.9}$$

where the last equality holds if $p(\mathbf{r}|s)$ is twice differentiable and the neural responses are conditionally independent (Lehmann and Casella, 1998). Assuming an unbiased estimator, the bound can be simplified to

$$\mathbb{E}[(G(s)-s)^2] = \mathrm{Var}(G(\mathbf{r})) \geq \frac{1}{J(s)}. \tag{S.10}$$

For multi-parameter estimation, let $J(\mathbf{s})$ denote the fisher information matrix, with elements defined analogous to Eq. S.9

$$J_{k,l}(\mathbf{s}) = -\mathbb{E}\left[\frac{\partial^2}{\partial s_k \partial s_l}\log p(\mathbf{r}|\mathbf{s})\right], \tag{S.11}$$

then (for unbiased estimators) the Cramér-Rao bound is instead stated as the following matrix inequality (Lehmann and Casella, 1998)

$$\mathrm{Cov}(G) = \Sigma \geq J^{-1}(\mathbf{s}) \tag{S.12}$$

in the sense that the difference $\Sigma - J^{-1}(\mathbf{s})$ is a positive semi-definite matrix. Thus, this implies the following lower bound for MSE of the k:th term

$$\mathrm{Var}(G^{(s_k)}) = \Sigma_{k,k} \geq \{J^{-1}(\mathbf{s})\}_{k,k} \geq \{J_{k,k}(\mathbf{s})\}^{-1} \tag{S.13}$$

where $G^{(s_k)} = \hat{s}_k$, i.e., the estimation of $s_k$ using estimator $G$. Note that the last inequality becomes an equality if $J(\mathbf{s})$ is a diagonal matrix, i.e., $\{J(\mathbf{s})\}_{j,k} = 0$ for all $j \neq k$.

For the tuning curves defined in Eq. S.1, the diagonal elements of the fisher information matrix can be analytically solved assuming $a_i \sim a$ within each module (see SI)

$$J_{k,k}(s) \approx (2\pi)^2 TN \frac{a}{R^2 w} B_0\left(\frac{1}{w}\right)^{D-1} \exp\left(-\frac{D}{w}\right) B_1\left(\frac{1}{w}\right)\overline{\lambda^{-2}} \tag{S.14}$$

where the bar indicates the average across modules. The off-diagonal elements, on the other hand, can be shown to be $0$ (see SI). Thus we have equality in the last inequality of Eq. S.13, and the MSE for each stimulus dimension is lower bounded by Eq. S.14.

#### 2.1 Approximating fisher information

To analytically approximate the fisher information for a given neural population, we will neglect the impact of spontaneous activity $b$. Then, the tuning curves in Eq. S.1 factorize as $f_i(s) = aq_{i,1}(s_1)...q_{i,D}(s_D)$ and the log-likelihood for $N$ neurons with conditionally independent spike counts becomes

$$\log p(\mathbf{r}|\mathbf{s}) = \sum_{i=1}^{N} r_i \log(Tf_i(\mathbf{s})) - Tf_i(\mathbf{s}) - \log r_i! \tag{S.15}$$

By taking the second derivatives w.r.t. stimulus dimension, we get for $k = l$:

$$\frac{\partial^2}{\partial s_k^2} \log p(\mathbf{r}|\mathbf{s}) = ... = \sum_{i}^{N} r\bigg(\frac{q''_{i,k}}{q_{i,k}} - \bigg(\frac{q'_{i,k}}{q_{i,k}}\bigg)^2\bigg) - Tf_i\frac{q''_{i,k}}{q_{i,k}} \tag{S.16}$$

and for $k \neq l$

$$\frac{\partial^2}{\partial s_k \partial s_l} \log p(\mathbf{r}|\mathbf{s}) = ... = \sum_{i}^{N} -Tf_i\frac{q'_k q'_l}{q_k q_l}. \tag{S.17}$$

Consequently, the elements of the fisher information matrix are given by

$$J_{k,k}(\mathbf{s}) = -\mathbb{E}\bigg[\frac{\partial^2}{\partial s_k^2} \log p(\mathbf{r}|\mathbf{s})\bigg] = \sum_{i=1}^{N} Tf_i(\mathbf{s})\bigg(\frac{q'_{i,k}(s_k)}{q_{i,k}(s_k)}\bigg)^2 \tag{S.18}$$

and for $k \neq l$

$$J_{k,l}(\mathbf{s}) = -\mathbb{E}\bigg[\frac{\partial^2}{\partial s_l \partial s_k} \log p(r|\mathbf{s})\bigg] = \sum_{i=1}^{N} -Tf_i(\mathbf{s})\frac{q'_k(s_k)q'_l(s_l)}{q_k(s_k)q_l(s_l)}. \tag{S.19}$$

To simplify calculations, it is possible to reparametrize the stimulus as in Eq. S.2 using the formula for fisher information under reparametrization (Lehmann and Casella, 1998)

$$J_{k,l}(\phi) = \sum_m \sum_n \frac{ds_m}{d\phi_k}\frac{ds_n}{d\phi_l} J_{k,l}(\mathbf{s}) = R^2 J_{k,l}(\mathbf{s}) \tag{S.20}$$

to obtain

$$J(\mathbf{s}) = \frac{1}{R^2} J_{k,l}(\phi). \tag{S.21}$$

We can approximate the elements of the fisher information matrix $J(\phi)$ in the limit of large $N$ by replacing the sums with integrals, e.g,

$$J_{k,k}(\phi) = \sum_{i=1}^{N} Tf_i(\phi)\bigg(\frac{q'_{i,k}(\phi_k)}{q_{i,k}(\phi_k)}\bigg)^2 \approx \tag{S.22}$$

$$\approx \sum_{j=1}^{L} \frac{L}{\lambda_j^D} a_j \int_{\phi_1 - \frac{1}{2}\lambda_j}^{\phi_1 + \frac{1}{2}\lambda_j} \cdots \int_{\phi_D - \frac{1}{2}\lambda_j}^{\phi_D + \frac{1}{2}\lambda_j} \bigg[\prod_{p=1}^{D} \exp\bigg(\frac{1}{w}\big(\cos(\frac{2\pi}{\lambda_j}(\phi_p - \phi'_p)) - 1\big)\bigg)\bigg] \frac{(2\pi)^2 \sin^2(\frac{2\pi}{\lambda_j}(\phi_k - \phi'_k))}{\lambda_j^2 w^2} d\phi' \tag{S.23}$$

where $L$ is the number of distinct modules, $M$ is the number of neurons in each module, $d\phi' = d\phi'_1...d\phi'_D$, and the D-dimensional integral is taken over the interval $[\phi_p - \frac{1}{2}\lambda_j, \phi_p + \frac{1}{2}\lambda_j)$ along each dimension. Making the variable substitution $\theta_p = \frac{2\pi}{\lambda_j}(\phi_p - \phi'_p)$ for $p = \{1, ..., D\}$ we have

$$J_{k,k}(\phi) \approx Ma \sum_{j=1}^{L} \frac{1}{\lambda_j^D} \int_{\pi}^{-\pi} \cdots \int_{\pi}^{-\pi} \bigg[\prod_{p=1}^{D} \exp\bigg(\frac{1}{w}\big(\cos(\theta_p) - 1\big)\bigg)\bigg] \frac{(2\pi)^2 \sin^2(\theta_k)}{\lambda_j^2 w^2} \frac{\lambda_j^D}{(-1)^D (2\pi)^D} d\theta = \tag{S.24}$$

$$= ... = \frac{(2\pi)^2 Na}{w} B_0\bigg(\frac{1}{w}\bigg)^{D-1} B_1\bigg(\frac{1}{w}\bigg) \exp\bigg(-\frac{D}{w}\bigg)\overline{\lambda^{-2}} \tag{S.25}$$

where the expectation is taken over the population's distribution of spatial frequencies and $B_\alpha(\cdot)$ is the modified Bessel function of the first kind. Similar calculations for the case $k \neq l$ yield

$$J_{k,l}(\phi) = ... \approx M \sum_{j=1}^{L} \frac{a}{w^2} \exp\Big(-\frac{D}{w}\Big) B_0\Big(\frac{1}{w}\Big)^{D-2} \int_{-\pi}^{\pi} \frac{1}{\lambda_j} \sin(\theta_k) \exp\Big(\frac{1}{w}\cos(\theta_k)\Big) d\theta_k \int_{-\pi}^{\pi} \frac{1}{\lambda_j} \sin(\theta_l) \exp\Big(\frac{1}{w}\cos(\theta_l)\Big) d\theta_l = \\ = 0. \tag{S.26}$$

Thus, the stimulus parameters will be asymptotically orthogonal for all of the populations considered in this paper. That is, the covariance matrix will be diagonal. The per-neuron average contribution to each diagonal element of the fisher information matrix, as reported in the main text, is therefore

$$\bar{J}_{k,k}(s) \approx \frac{(2\pi)^2 a}{R^2 w} B_0\Big(\frac{1}{w}\Big)^{D-1} B_1\Big(\frac{1}{w}\Big) \exp\Big(-\frac{D}{w}\Big) \overline{\lambda^{-2}}. \tag{S.27}$$

### 3 Calculate $M_{k,l,m}$

We will need to calculate $\frac{\partial^3}{\partial s_m \partial s_l \partial s_k} \log p(\mathbf{r}|\mathbf{s})$. For $k \neq l \neq m$ we have,

$$\frac{\partial^3}{\partial s_m \partial s_l \partial s_k} \log p(\mathbf{r}|\mathbf{s}) = -\sum_{i=1}^{N} T f_i(\mathbf{s}) \frac{q_k'(s_k) q_l'(s_l) q_m'(s_m)}{q_k(s_k) q_l(s_l) q_m(s_m)} \tag{S.28}$$

Thus, $M_{k,l,m}$ for $k \neq l \neq m$ becomes

$$M_{k,l,m} = -\sum_{i=1}^{N} T f_i(\mathbf{s}^*) \frac{q_k'(s_k^*) q_l'(s_l^*) q_m'(s_m^*)}{q_k(s_k^*) q_l(s_l^*) q_m(s_m^*)} \approx \tag{S.29}$$

$$\approx \sum_{j=1}^{L} \frac{M}{\lambda_j^D} \int_{s_1^*-\frac{R}{2}\lambda_j}^{s_1^*+\frac{R}{2}\lambda_j} \cdots \int_{s_D^*-\frac{R}{2}\lambda_j}^{s_D^*+\frac{R}{2}\lambda_j} Ta\Big[\prod_{j=1}^{D} \exp\Big(\frac{1}{w}\Big(\cos\big(\frac{2\pi}{\lambda_j R}(s_j^* - s_j')\big) - 1\Big)\Big)\Big] \frac{(2\pi)^3}{\lambda_j^3 w^3} \sin\Big(\frac{2\pi}{\lambda_j R}(s_k^* - s_k')\Big) \tag{S.30}$$

$$\sin\Big(\frac{2\pi}{\lambda_j R}(s_l^* - s_l')\Big) \sin\Big(\frac{2\pi}{\lambda_j R}(s_m^* - s_m')\Big) d\mathbf{s}' = 0 \tag{S.31}$$

as odd functions over even intervals integrate to zero. For $k \neq l = m$ (note that $k = l \neq m$ and $k = m \neq l$ follows by symmetry) we have

$$\frac{\partial^3}{\partial s_l^2 \partial s_k} \log p(\mathbf{r}|\mathbf{s}) = -\sum_{i=1}^{N} T f_i(\mathbf{s}) \frac{q_k'(s_k) q_l''(s_l)}{q_k(s_k) q_l(s_l)} \tag{S.32}$$

and hence,

$$M_{k,l,l} = -\sum_{i=1}^{N} T f_i(\mathbf{s}^*) \frac{q_k'(s_k^*) q_l''(s_l^*)}{q_k(s_k^*) q_l(s_l^*)} \approx \\ \approx -\sum_{j=1}^{L} \frac{M}{\lambda_j^D} \int_{s_1^*-\frac{R}{2}\lambda_j}^{s_1^*+\frac{R}{2}\lambda_j} \cdots \int_{s_D^*-\frac{R}{2}\lambda_j}^{s_D^*+\frac{R}{2}\lambda_j} Ta\Big[\prod_{j=1}^{D} \exp\Big(\frac{1}{w}\Big(\cos\big(\frac{2\pi}{\lambda_j R}(s_j^* - s_j')\big) - 1\Big)\Big)\Big] \frac{(2\pi)^3}{\lambda_j^3 w^3} \sin\Big(\frac{2\pi}{\lambda_j R}(s_k^* - s_k')\Big) \\ \Big(w \cos\Big(\frac{2\pi}{\lambda_j R}(s_l^* - s_l')\Big) - \sin^2\Big(\frac{2\pi}{\lambda_j R}(s_l^* - s_l')\Big)\Big) d\mathbf{s}' = 0. \tag{S.33}$$

Lastly, for $k = l = m$ we have,

$$\frac{d^3}{ds_k^3} \log p(\mathbf{r}|\mathbf{s}) = \sum_{i=1}^{N} (r_i - T f_i(\mathbf{s})) \frac{q_k'''(s_k)}{q_k(s_k)} - 3 r_i \frac{q_k'(s_k) q_k''(s_k)}{q_k(s_k)^2} + 2 r_i \Big(\frac{q_k'(s_k)}{q_k(s_k)}\Big)^3 \tag{S.34}$$

thus, $M_{k,k,k}$ becomes

$$M_{k,k,k} = \mathbb{E}_{\mathbf{r}\sim\text{Poiss}(f(\mathbf{s}^\circ))}\left[\frac{d^3}{ds_k^3}\log p(\mathbf{r}|\mathbf{s}^*)\right] = \sum_{i=1}^{N}(Tf_i(\mathbf{s}^\circ) - Tf_i(\mathbf{s}^*))\frac{q_k'''(s_k^*)}{q_k(s_k^*)} - 3Tf_i(\mathbf{s}^\circ)\frac{q_k'(s_k^*)q_k''(s_k^*)}{q_k(s_k^*)^2} + 2Tf_i(\mathbf{s}^\circ)\left(\frac{q_k'(s_k^*)}{q_k(s_k^*)}\right)^3 \tag{S.35}$$

We approximate this sum by an $D$-dimensional integral, let $f_{(j)}(s,s')$ denote any tuning curve with $\lambda = \lambda_j$ and firing field given by $s' = (s_1', ..., s_D')$

$$M_{k,k,k} = \mathbb{E}_{\mathbf{r}\sim\text{Poiss}(f(\mathbf{s}^\circ))}\left[\frac{d^3}{ds_k^3}\log p(\mathbf{r}|\mathbf{s}^*)\right] = \sum_{i=1}^{N}(Tf_i(\mathbf{s}^\circ) - Tf_i(\mathbf{s}^*))\frac{q_k'''(s_k^*)}{q_k(s_k^*)} - 3Tf_i(\mathbf{s}^\circ)\frac{q_k'(s_k^*)q_k''(s_k^*)}{q_k(s_k^*)^2} + 2Tf_i(\mathbf{s}^\circ)\left(\frac{q_k'(s_k^*)}{q_k(s_k^*)}\right)^3 \tag{S.36}$$

$$\approx T\sum_{j=1}^{L}\frac{M}{(\lambda_j R)^D}\int\frac{(2\pi)^3}{\lambda_j^3 R^3 w}\Bigg(-\frac{\sin^3\left(\frac{2\pi(s_k^*-s_k')}{\lambda_j R}\right)}{w^2} + 3\frac{\sin\left(\frac{2\pi(s_k^*-s_k')}{\lambda_j R}\right)\cos\left(\frac{2\pi(s_k^*-s_k')}{\lambda_j R}\right)}{w} + \tag{S.37}$$

$$+\sin\left(\frac{2\pi(s_k^*-s_k')}{\lambda_j R}\right)\Bigg)\left[f_{(j)}(\mathbf{s}^\circ,\mathbf{s}') - f_{(j)}(\mathbf{s}^*,\mathbf{s}')\right] + 3\frac{(2\pi)^3}{\lambda_j^3 R^3 w^2}\Bigg(\frac{\sin^3\left(\frac{2\pi(s_k^*-s_k')}{\lambda_j R}\right)}{w} + \tag{S.38}$$

$$-\sin\left(\frac{2\pi(s_k^*-s_k')}{\lambda_j R}\right)\cos\left(\frac{2\pi(s_k^*-s_k')}{\lambda_j R}\right)\Bigg)f_{(j)}(\mathbf{s}^\circ,\mathbf{s}') - 2\frac{(2\pi)^3}{\lambda_j^3 R^3 w^3}\sin^3\left(\frac{2\pi(s_k^*-s_k')}{\lambda_j R}\right)f_{(j)}(\mathbf{s}^\circ,\mathbf{s}')ds' = \tag{S.39}$$

$$= \left\{\text{simplifying by pairing all } f_{(j)}(s^\circ,s') \text{ and all } f_{(j)}(s^*,s') \text{ gives}\right\} = \tag{S.40}$$

$$= T\sum_{j=1}^{L}\frac{M}{(\lambda_j R)^D}\frac{(2\pi)^3}{\lambda_j^3 R^3 w}\int\sin\left(\frac{2\pi(s_k^*-s_k')}{\lambda_j R}\right)f_{(j)}(s^\circ,s') + \Bigg(\frac{\sin^3\left(\frac{2\pi(s_k^*-s_k')}{\lambda_j R}\right)}{w^2} - 3\frac{\sin\left(\frac{2\pi(s_k^*-s_k')}{\lambda_j R}\right)\cos\left(\frac{2\pi(s_k^*-s_k')}{\lambda_j R}\right)}{w} + \tag{S.41}$$

$$-\sin\left(\frac{2\pi(s_k^*-s_k')}{\lambda_j R}\right)\Bigg)f_{(j)}(s^*,s')ds' \tag{S.42}$$

Using the variable substitution $\theta_p^* = \frac{2\pi\xi_j}{R}(s_p^* - s_p')$, $p = \{1, ..., D\}$

$$M_{k,k,k} \approx T\sum_{j=1}^{L}\frac{M}{(\lambda_j R)^D}\frac{(2\pi)^3}{\lambda_j^3 R^3 w}a\int_{-\pi}^{\pi}\cdots\int_{-\pi}^{\pi}\sin\left(\theta_k^*\right)\left[\prod_{p=1}^{D}\exp\left(\frac{1}{w}\left(\cos\big(\overbrace{\frac{2\pi}{\lambda_j R}(s_p^\circ - s_p^*)}^{=\psi_p(\lambda_j)}+\theta_p^*\big) - 1\right)\right)\right] + \tag{S.43}$$

$$+\underbrace{\left(\frac{\sin^3(\theta_k^*)}{w^2} - 3\frac{\sin(\theta_k^*)\cos(\theta_k^*)}{w} - \sin(\theta_k^*)\right)\left[\prod_{p=1}^{D}\exp\left(\frac{1}{w}\left(\cos(\theta_p^*) - 1\right)\right)\right]}_{\text{odd function over even interval }=0}\frac{\lambda_j^D R^D}{(2\pi)^D}d\theta^* = \tag{S.44}$$

$$= T\sum_{j=1}^{L}\frac{M}{(2\pi)^D}\frac{(2\pi)^3 a}{\lambda_j^3 R^3 w}\int_{-\pi}^{\pi}\cdots\int_{-\pi}^{\pi}\sin\left(\theta_k^*\right)\left[\prod_{p=1}^{D}\exp\left(\frac{1}{w}\left(\cos\left(\psi_p(\lambda_j) + \theta_p^*\right) - 1\right)\right)\right]d\theta^* = \tag{S.45}$$

$$= T\sum_{j=1}^{L}\frac{M}{(2\pi)^D}\frac{(2\pi)^3 a}{\lambda_j^3 R^3 w}\exp\left(-\frac{D}{w}\right)\int_{-\pi}^{\pi}\cdots\int_{-\pi}^{\pi}\sin\left(\theta_k^*\right)\left[\prod_{p=1}^{D}\exp\left(\frac{1}{w}\cos\left(\psi_p(\lambda_j) + \theta_p^*\right)\right)\right]d\theta^* = \tag{S.46}$$

$$= T\sum_{j=1}^{L}\frac{M}{(2\pi)^D}\frac{(2\pi)^3 a}{\lambda_j^3 R^3 w}\exp\left(-\frac{D}{w}\right)\int_{-\pi}^{\pi}\sin\left(\theta_k^*\right)\exp\left(\frac{1}{w}\cos\left(\psi_k(\lambda_j) + \theta_k^*\right)\right)d\theta_k^*\left[\prod_{p\neq k}\int_{-\pi}^{\pi}\exp\left(\frac{1}{w}\cos\left(\psi_p(\lambda_j) + \theta_p^*\right)\right)d\theta_p^*\right] \tag{S.47}$$

$$= T\sum_{j=1}^{L}M\frac{(2\pi)^2 a}{\lambda_j^3 R^3 w}\exp\left(-\frac{D}{w}\right)B_0\left(\frac{1}{w}\right)^{(D-1)}\int_{-\pi}^{\pi}\sin\left(\theta_k^*\right)\exp\left(\frac{1}{w}\cos\left(\psi_k(\lambda_j) + \theta_k^*\right)\right)d\theta_k^* \tag{S.48}$$

Because $s^*$ depends on $r$, which will be different for every trial, so does $\theta^*$. We instead focus on the $\theta^*$ which maximizes the above integrals. Unfortunately, although this allows us to choose $\theta^*$, for each $\theta^*$ also we have a collection of $\psi_p(\lambda_j)$ as these variables also depend on $\lambda_j$. To obtain an analytical solution, we upper bound the above equation using the individual maximums for $\psi_p(\lambda_j)$. These terms then becomes independent of $\lambda_j$ but not on $w$, the validity of this approximation should decrease with smaller tuning widths.

Using these choices of $\psi_p(\lambda_j)$, we can calculate an upper bound to the maximum value of $M_{k,k,k}$ as

$$\max_{s^*} M_{k,k,k} \leq M^* = T\sum_{j=1}^{L} M\frac{(2\pi)^2 a}{\lambda_j^3 R^3 w}\exp\Big(-\frac{D}{w}\Big)B_0\Big(\frac{1}{w}\Big)^{(D-1)}\underbrace{\int_{-\pi}^{\pi}\sin\Big(\theta_k^*\Big)\exp\Big(\frac{1}{w}\cos\big(\psi^*+\theta_k^*\big)\Big)d\theta_k^*}_{C^*(w)} = \tag{S.49}$$

$$= TN\frac{(2\pi)^2 a}{R^3 w}\exp\Big(-\frac{D}{w}\Big)B_0\Big(\frac{1}{w}\Big)^{(D-1)}C^*(w)\mathbb{E}\Big[\lambda^{-3}\Big] \tag{S.50}$$

### 4 Approximating minimal required spike count

Given the approximation of minimal decoding time in Eq. 47, we seek to reformulate the approximation in terms of required total spike count, instead. The average total spike count for a given population and stimulus condition is

$$\mu(\mathbf{s}) = \mathbb{E}_r\Big[\sum_{i=1}^{N} r_i\Big|\mathbf{s}\Big] = \sum_{i=1}^{N} Tf_i(\mathbf{s}) \tag{S.51}$$

where $T$ is the decoding time. Thus, the average spike count over both stimulus conditions (assuming uniformly distributed stimulus) and trials is

$$\mu = \mathbb{E}_{\mathbf{s}}\Big[\mathbb{E}_{\mathbf{r}}\Big[\sum_{i=1}^{N} r_i\Big|\mathbf{s}\Big]\Big] = \frac{1}{R^D}\int_0^R\cdots\int_0^R\sum_{i=1}^{N} Tf_i(\mathbf{s})d\mathbf{s} = NT(aB_0(1/w)^D\exp(-D/w)+b). \tag{S.52}$$

Consequently, the number of spikes evoked by the stimulus-related tuning of the population is

$$\mu_{stim} = NTaB_0(1/w)^D\exp(-D/w). \tag{S.53}$$

Inserting Eq. S.53 into Eq. 46 reveals the number of stimulus-evoked spikes, $\mu^*_{stim}$, the population must produce before reaching the theoretical lower bound

$$\mu^*_{stim} \approx K_1^* A(w)B_0(1/w)\frac{\overline{\lambda^{-3}}^2}{\overline{\lambda^{-2}}^3} + K_2^*. \tag{S.54}$$

##### 4.1 Minimal required spike count in simulation

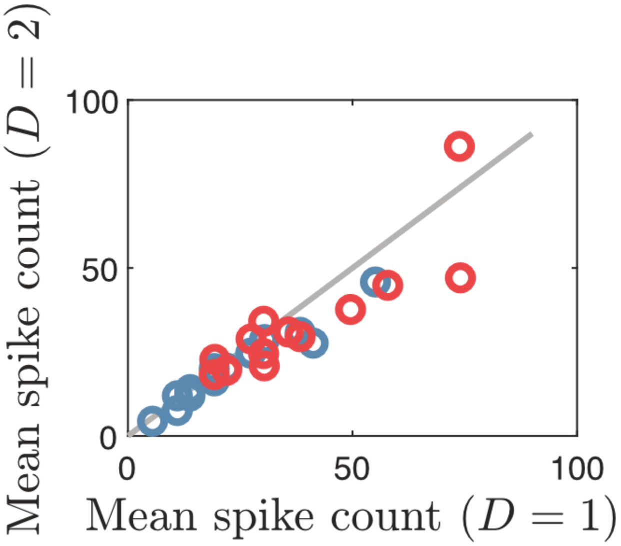


Figure S1: Mean spike count required to remove catastrophic errors for populations in Fig. 2. Each circle indicate the minimal spike count for a single population with constant scale factor but encoding either a 1-dimensional (x-axis) or a 2-dimensional stimulus (y-axis). Blue circles indicate $\lambda_1 = 1$ and red circles $\lambda_1 = 1/2$. Being on the grey line corresponds to having the same required spike count for both stimulus cases.

#### 5 Comparing actual stimulus estimation

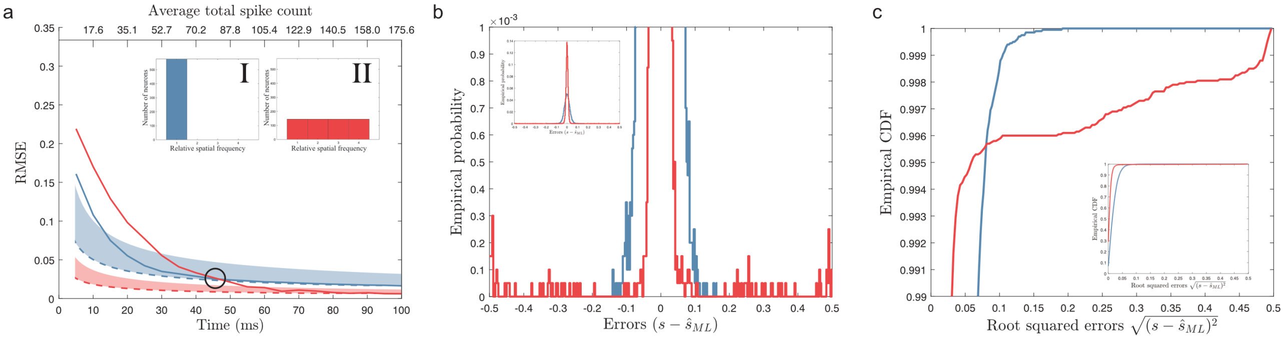


Figure S2: Comparing actual stimulus estimation. a) The time evolution of the RMSE (solid lines) of two populations indicated by $\mathbf{I}$ and $\mathbf{II}$. Dashed lines indicate the lower bound predicted by Cramér-Rao and the shaded areas indicate the region between the lower bound and the limit set for minimal decoding times. The parameters were set according to Table. S3 using $b = 2$ sp/s and $D = 2$. The black circle indicates the point where population $\mathbf{I}$ and $\mathbf{II}$ have roughly equal RMSE. Note that relative spatial frequency is the inverse of relative spatial period, i.e., $1/\lambda$. b) Plotting the empirical probability for the time point where population $\mathbf{I}$ and $\mathbf{II}$ have roughly equal RMSE (or MSE) reveals the problem of describing errors using RMSE. Although the RMSE is roughly equal, the distribution of errors are highly different. Inset: While population $\mathbf{I}$ has a relatively wide distribution of errors centered around $0$, the population $\mathbf{II}$ has a much more narrow distribution. Full plot: zooming in on very rate error events (probability $0 - 0.1\%$) reveals that while the multi-peaked population has a narrower distribution of errors around $0$, it also has almost uniform distribution of rare errors ranging over the entire stimulus domain. c) The empirical CDF of the root squared errors for the same two populations as in b) highlights the heavy tailed distribution of errors for the multi-peaked population $\mathbf{II}$. Inset: empirical CDF across all root squared errors.

To exemplify the issue with MSE as a descriptor of the error distribution, we compared the time evolution of root mean squared errors (RMSEs) for two different populations encoding a two-dimensional stimulus $(D = 2)$. Note that the choice of RMSE does not qualitatively change any results, but only allows the errors to be compared to the normalized stimulus range. This time, however, we evaluated the performance based on 10000 predefined stimulus conditions $(D = 2)$ that were randomly and independently sampled from a uniform distribution. These stimulus conditions were then reused for both populations and all choices of decoding time. The time evolution of the populations' RMSEs are plotted in Fig. S2a. As expected, the single-peaked population, $\mathbf{I}$, is better suited to reduce RMSE at very short time scales. However, after around $45$ ms or roughly 80 spikes, the periodic population, $\mathbf{II}$, has a lower RMSE. Comparing the distributions of errors for the decoding time when population $\mathbf{I}$ and $\mathbf{II}$ are roughly equal (indicated by black circle in Fig. S2a) reveals a more narrow distribution of errors for the periodic population $\mathbf{II}$ (Fig. S2b). This is predicted by the asymptotically normal distribution of local errors being smaller for the periodic population (Lehmann and Casella, 1998). However, in contrast to population $\mathbf{I}$, population $\mathbf{II}$ also has occasional errors spanning the entire stimulus domain (see Fig. S2b). These large errors are unlikely to come from the normal distribution of local errors and are therefore, intuitively, catastrophic errors.

The differences in the distributions of errors between population $\mathbf{I}$ and $\mathbf{II}$ are more clearly visible when looking at the cumulative distribution of root squared errors (Fig. S2c). While a larger fraction of errors are close to $0$ for population $\mathbf{II}$, the last percent of errors spans the entire stimulus domain. In comparison, population $\mathbf{I}$ has a wider distribution of local errors but a sharp threshold on the maximal error which is much smaller than the maximal error for population $\mathbf{II}$. This suggests that although the MSEs are roughly equal, the distributions of errors are very different. While population $\mathbf{II}$ has smaller local errors than population $\mathbf{I}$, it still experiences catastrophic errors.

As is demonstrated in Fig. S2, comparing the appropriateness of the neural code solely based on MSE can be highly misleading. Different population can have similar MSE yet have different distributions of errors. Thus, if one accepts the premise that catastrophic estimation errors, although rare, should be especially detrimental for sensory processing, then MSE and fisher information are only useful descriptors of the accuracy when coupled to the minimal decoding time.

### 6 Tables

| Parameters | Parameter values |
|---|---|
| Number of neurons, N | $600$ |
| Number of modules, $L$ | $2$ |
| Scale factor, $c$ | 0.05 - 1 |
| Average firing rate (sp/s) | $20\exp(-1/w)B_0(1/w)$ |
| Peak amplitude, $a_i$ | $\frac{20\exp(-1/w)B_0(1/w)}{\int_0^R q_i(s)ds}$ |
| Width parameter, $w$ | 0.3 |
| Spontaneous activity, $b$ (spikes/s) | 0 |

Table S1: Parameters and parameter values for neural populations in Fig. 1.

| Parameters | Parameter values |
|---|---|
| Number of neurons, N | $600$ |
| Number of modules, $L$ | $5$ |
| Scale factor, $c$ | 0.3 - 1 |
| Average firing rate (sp/s) | $20\exp(-D/w)B_0(1/w)^D$ |
| Peak amplitude, $a_i$ | $\frac{20\exp(-1/w)B_0(1/w)}{\int_0^R q_i(s)ds}$ |
| Width parameter, $w$ | 0.3 |
| Spontaneous activity, $b$ (spikes/s) | 0 (Fig. 2) or 2 (Fig. 3) |

Table S2: Parameters and parameter values for neural populations in Fig. 2 and 3.

| Parameters | Parameter values |
|---|---|
| Number of neurons, N | $12^2 = 576$ |
| Amplitudes, $a$ (spikes/s) | 20 |
| Spatial frequencies | $[1]$, $[1, 2, 3, 4]$ |
| Width parameter, $w$ | 0.3 |
| Spontaneous activity, $b$ (spikes/s) | 2 |

Table S3: Parameters and parameter values for neural populations for Fig. S2.

| Parameters | Parameter values |
|---|---|
| $\tau_s$ (s) | 0.5 |
| $\sigma_s$ | 0.1 |

Table S4: Parameters and parameter values for OU stimulus.

| Parameters | Parameter values |
|---|---|
| Membrane time constant, $\tau_{memb}$ (ms) | $20$ |
| Threshold memb. potential, $V_{th}$ (mV) | $20$ |
| Reset memb. potential (mV) | $10$ |
| Resting potential, $V_0$ (mV) | $0$ |
| Refractory period, $\tau_{rp}$ (ms) | $2$ |

Table S5: Parameters and parameter values for LIF neurons.

| Parameters | Parameter values |
|---|---|
| Number of neurons 1st layer, $N_1$ | $500$ |
| Number of neurons 2nd layer, $N_2$ | $400$ |
| Maximal stimulus-dependent input rate, $a$ (spikes/s) | $750$ |
| Baseline input rate, $b$ (spikes/s) | $4250$ |
| Relative spatial periods, $\lambda_j$ | $[1]$ or $[1, 2, 3, 4]$ |
| Width parameter, $w$ | $0.3$ |
| Width parameter, $w_{ro}$ | $\frac{(\pi/N_2)^2}{2\log(2)}$ |
| Input EPSP (1st layer), $J_E$ (mV) | $0.2$ |
| Maximal EPSP (2nd layer), $J_{EE}$ (mV) | $2$ |
| Maximal IPSP (2nd layer), $J_{II}$ (mV) | $2$ |
| Synaptic delays, $d$ (ms) | $1.5$ |

Table S6: Spiking network parameters and parameter values.

### 7 Pseudo-codes

These pseudo-codes are mean as a guidance to make both the code and the results more easy to interpret.

**Algorithm 1** Estimate MSE

**Require:** $\mathbf{f}(s), T, D$

$counterLimit \leftarrow 1'000$ ▷ # of consecutive estimated samples with stable MSE before exit
$MSE^{(old)} \leftarrow 0$
$counter \leftarrow 0$ ▷ counter for # consecutively estimated samples with stable MSE
**while** $counter < counterLimit$ **do**
  $\mathbf{s} \sim$ Uniform$([0,1)^D)$ ▷ Sample a stimulus condition
  $\mathbf{r} \sim \Big(\text{Poission}\big(Tf_1(s)\big), ..., \text{Poission}\big(Tf_N(s)\big)\Big)$ ▷ Sample spike counts from Poisson distributions
  $\hat{s}_{ML} \leftarrow$ MLE$(\mathbf{s}, \mathbf{r})$ ▷ Estimate the stimulus using maximum likelihood estimation
  $MSE \leftarrow$ UpdateMSE$(\hat{s}_{ML}, MSE^{(old)})$
  $d \leftarrow$ getNonZeroDecimals$(MSE)$
  **if** floor$(10^d MSE) ==$ floor$(10^d MSE^{(old)})$ **then**
    $counter \leftarrow counter + 1$ ▷ If MSE up to the first 2 non-zero decimals remains constant, add one to the counter
  **else**
    $counter \leftarrow 0$ ▷ If MSE up to the first 2 sig. decimals changed, reset counter
  **end if**
  $MSE^{(old)} \leftarrow MSE$
**end while**
**return** $MSE$

**Algorithm 2** Minimal decoding time

**Require:** $\mathbf{f}(s), D$ ▷ $\mathbf{f}(s)$ = the tuning curves of the population, $D$ = the stimulus dimensionality

$T \leftarrow 0$ ▷ $T$ decoding time
**while** $continue == true$ **do**
  $T \leftarrow T + 10^{-3}$
  $J \leftarrow$ getFI$(\mathbf{f}(s), T, D)$ ▷ Calculate Fisher information using decoding time $T$
  $MSE \leftarrow$ estimateMSE$(f, T, D)$ ▷ Using the algorithm sketched above
  **if** mean$(MSE) \leq 2 \cdot$ mean(diag$(J^{-1})$) **then**
    $continue \leftarrow false$ ▷ If empirical MSE within 2 times the lower bound -> exit
  **end if**
**end while**
**return** $T$
